# CryoRhodopsins: a comprehensive characterization of a group of microbial rhodopsins from cold environments

**DOI:** 10.1101/2024.01.15.575777

**Authors:** G.H.U. Lamm, E. Marin, A. Alekseev, A.V. Schellbach, A. Stetsenko, G. Bourenkov, V. Borshchevskiy, M. Asido, M. Agthe, S. Engilberge, S.L. Rose, N. Caramello, A. Royant, T. R. Schneider, A. Bateman, T. Mager, T. Moser, J. Wachtveitl, A. Guskov, K. Kovalev

## Abstract

Microbial rhodopsins are omnipresent on Earth, however the vast majority of them remain uncharacterized. Here we describe a new rhodopsin group from cold-adapted organisms and cold environments, such as glaciers, denoted as CryoRhodopsins (CryoRs). Our data suggest that CryoRs have dual functionality switching between inward transmembrane proton translocation and photosensory activity, both of which can be modulated with UV light. CryoR1 exhibits two subpopulations in the ground state, which upon light activation lead to transient photocurrents of opposing polarities. A distinguishing feature of the group is the presence of a buried arginine residue close to the cytoplasmic face of its members. Combining single-particle cryo-electron microscopy and X-ray crystallography with the rhodopsin activation by light, we demonstrate that the arginine stabilizes a UV-absorbing intermediate of an extremely slow CryoRhodopsin photocycle. Together with extensive spectroscopic characterization, our investigations on CryoR1 and CryoR2 proteins reveal mechanisms of photoswitching in the newly identified group and demonstrate principles of the adaptation of these rhodopsins to low temperatures.

## INTRODUCTION

Microbial rhodopsins (MRs) constitute a large family of light-sensitive membrane proteins found all over the Earth in various organisms such as Archaea, Bacteria, Eukaryota, and Viruses^1,2^. Due to the continuous sampling and metagenomics analysis the phylogenetic tree of MRs is constantly expanding^1,3^. This expansion affects both the known groups when more of their members are being identified and assigned to these groups but also the discovery of novel clades of MRs, which may form distinct branches in the phylogenetic tree^1,4^. The characterization of novel MRs is crucial for fundamental science as it opens the door for revealing new molecular mechanisms of ion transport, light sensing, signal transduction, enzymatic activities, etc. Moreover, beyond the ecological, evolutionary, biochemical, and biophysical relevance it is of great interest and importance also for the biotechnological applications of MRs such as optogenetics^5^. For instance, recently discovered light-gated anion and cation channelrhodopsins already entered use as highly effective optogenetic tools^6–8^.

One approach for revealing novel groups of MRs uses bioinformatics search within the gene databases and sequence analysis to find proteins with unusual functional motifs or domain architecture. Typically, 3- or 6-letter motifs are analyzed, which include the residues in helices C and G at the positions of D85, T89, D96 or R82, D85, T89, D96, D212, and K216 in the classical representative of MRs, bacteriorhodopsin from *Halobactrium salinarum* (BR)^9^. However, recent works showed that there are other positions in MRs, which co-determine the function of the protein and might differ between the groups of a given family. An example is the position of T46 in the helix B of BR, which can be occupied by glutamic acid in viral rhodopsins of groups 1 and 2^10,11^ or histidine in bacterial outward proton pumps^12,13^ and is actively involved in the ion translocation process.

In this work, we used bioinformatics and found MRs with unusual 7-letter motifs (corresponding to the residues T46, R82, D85, T89, D96, D212, and K216 of BR), which can be assigned to a separate group in the phylogenetic tree. This group, characterized by the presence of an arginine residue at the position of T46 in BR, has never been described before to our knowledge. The members of the group were restricted to cold environments and were found in cold-living bacteria, such as *Cryobacterium*, *Glacier habitats*, *Subtercola sp*., and others. Therefore, we named this group as CryoRhodopsins (CryoRs). We show that CryoRs have extremely slow photocycles dominated by the blue-shifted intermediate state. High-resolution single-particle cryo-electron microscopy (cryo-EM) and X-ray crystallography on two CryoRs demonstrated three key roles of the characteristic arginine in helix B in the stabilization of this intermediate. At the same time, we demonstrate that CryoRs vary in their spectral properties. For instance, some of them exhibited strong pH dependence of the absorption spectra and photocycle kinetics, while in others it is negligible. Strikingly, one rhodopsin (CryoR1 from *Cryobacterium levicorallinum*) shifts its absorption maximum in a wide range of pH and shows an 80 nm red shift to 620 nm below pH 6. Furthermore, CryoR1 displays strong coherent oscillations on the ultrafast timescale in its 620 nm absorbing state at low pH indicating a hindered isomerization of the retinal chromophore. Electrophysiology of CryoR1 revealed that this 620 nm absorbing state likely corresponds to a light-driven inward proton-pumping modality of the protein. Thus, our findings suggest dual functionality of CryoRs as either photosensors at neutral and alkaline pH, or inward proton pumps at low pH. We also demonstrate the use of cryo-EM for the determination of high-resolution structures of small membrane proteins such as rhodopsins in the intermediate states following light activation to uncover the molecular mechanisms of the CryoRs work.

## RESULTS

### Identification of CryoRhodopsins

To identify new unusual groups of MRs we performed a bioinformatics search within the UniProtKB, UniParc, Genbank, and MGnify gene databases and analyzed the 7-letter motifs of the output protein sequences. As a result, we found MRs with the motif pattern RRXXXDK, where XXX varied between EAE, ESE, ETE, DSE, DNE, DTE, GSE, or GTQ. In total, we identified 40 unique complete protein sequences with these functional motifs, which clustered together in the phylogenetic tree (Fig. 1A,B).

**Fig. 1.**
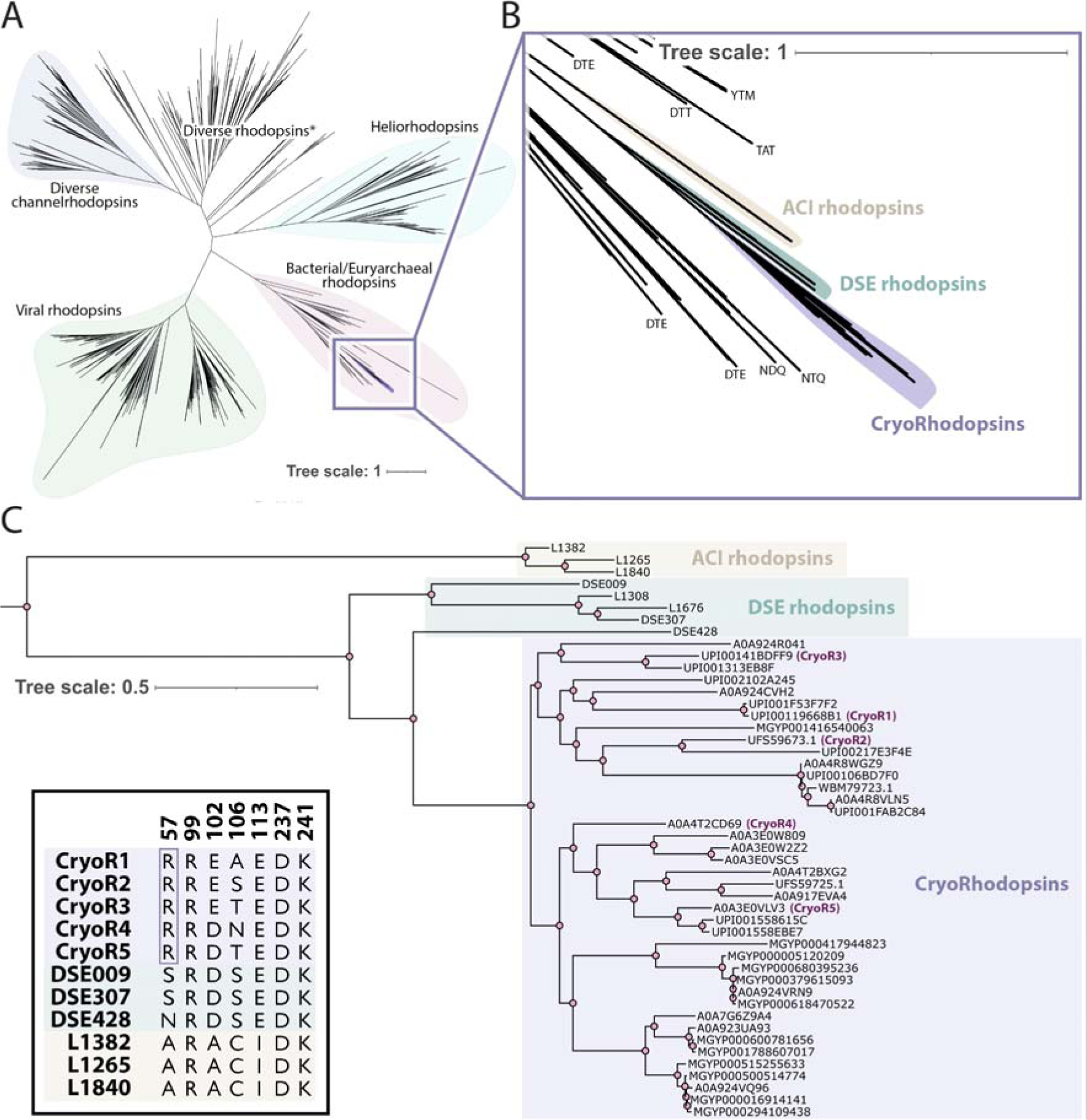
Phylogeny of CryoRhodopsins. **A.** Maximum likelihood phylogenetic tree of MRs. The tree includes 2199 sequences reported in ref. ^1^, 3 sequences of DSE rhodopsins reported in ref. ^14^, and 40 sequences of CryoRhodopsins found in the present work. **B.** Enlarged view of the tree branch containing CryoRs. Characteristic 3-letter motifs of the nearby branches are shown at the tips. **C.** Rectangular representation of the phylogenetic tree of the CryoRs and nearby DSE and ACI rhodopsins groups. The inset in the left bottom corner shows amino acids of the 7-letter motifs of CryoR1-5, DSE, and ACI rhodopsins (numbering corresponds to CryoR1). The unique arginine (R57 in CryoR1) is boxed for clarity.

Surprisingly, the members of this new group of MRs were found only in cold environments. Specifically, 9 and 12 rhodopsins of a total of 29 genes with identified origin belong to genera *Cryobacterium* and *Subtercola*, respectively (Table S1). Since the newly discovered rhodopsins come from cold environments and are well-distributed in 9 various species of *Cryobacterium* (versus 4 different species of *Subtercola*), we named this separate group CryoRhodopsins (CryoRs) to reflect both their origin and the link to areas with low temperatures.

Phylogenetically, CryoRs are located closely to the branch of the bacterial proton, sodium, and chloride pumps (Fig. 1A,B). It should be noted that rhodopsins with EAE motif appeared in the phylogenetic tree in ref. ^15^ and were assigned to the P1 family of MRs together with the recently reported group of DSE rhodopsins^14^ and some other previously-studied proteins with DTE motif^16–18^. The closest relatives of CryoRs are DSE rhodopsins^14^ (Fig. 1B,C). The analysis of the protein sequences showed that both CryoRs and DSE rhodopsins have common features such as elongated N- and C-termini (Fig. S1); however DSE rhodopsins do not possess the arginine found in the helix B of CryoRs (Fig. 1C). Moreover, DSE rhodopsins do not originate from cold environments. Thus, we conclude that the arginine residue is a unique characteristic feature of CryoRs. Indeed, no MRs with an arginine residue at the position of T46 in BR have been reported. The next closest relatives are rhodopsins with unusual ACI motif^1^, which differ significantly from both the DSE rhodopsins and CryoRs (Fig. 1B,C).

The majority of the CryoRs (39 of 40) possesses a glutamic acid residue at the position corresponding to that of a primary proton donor to the retinal Schiff base (RSB), D96 in BR^19^. This position is occupied by a glutamic acid acting as a proton donor in most light-driven bacterial outward proton pumps^3^. In contrast to the arginine and glutamic acid at the cytoplasmic side, the set of two key residues of the functional motif located closely to the RSB in the central region of the rhodopsin varies largely within the group. Nevertheless, 38 out of 40 CryoRs have a carboxylic residue (15 - glutamic acid; 23 - aspartic acid) at the position of a primary proton acceptor D85 in BR^20^. Thus, most CryoRs have two RSB counterions, similar to BR and many other MRs. The amino acid residue at the position of T89 in BR is occupied by alanine (8), serine (7), threonine (24), or asparagine (1).

For the in-depth investigations, we selected 5 representatives of the CryoR group with different 7-letter motifs: RREAEDK (CryoR1, *Cryobacterium levicorallinum*, GenBank ID: WP_166787544), RRESEDK (CryoR2, *Subtercola sp*. AK-R2A1-2, GenBank ID: UFS59673.1), RRETEDK (CryoR3, *Cryobacterium frigoriphilum*, GenBank ID: WP_166791776), RRDNEDK (CryoR4, *Subtercola vilae*, UniProt ID: A0A4T2CD69), and RRDTEDK (CryoR5, *Subtercola boreus*, UniProt ID: A0A3E0VLV3) (Fig. 1C). Below, we describe the common spectral and structural features of CryoRs originating from the presence of the arginine residue at their cytoplasmic side as well as the differences between the CryoRs and their new unique properties.

### Slow photocycle kinetics of CryoRs dominated by blue-shifted intermediate

CryoR1-5 differed in their UV-Vis absorption spectra with the main band position varying in the range of 390-580 nm (Fig. 2; Fig. S2A). However, the studied proteins have a common feature in their spectral behavior, which is the presence of a long-living blue-shifted intermediate state of the photocycle (Fig. 2). The intermediate lasts for minutes and accumulates in the samples upon illumination as indicated by a color change of all studied CryoRs when exposed to ambient light in the laboratory. To investigate such photoconversion of CryoRs in more detail, we performed illumination experiments of CryoR1-5 using LEDs with central wavelengths corresponding to the main absorption band in the green-to-red spectral area. Due to the unusual sensitivity of CryoRs to light, the samples were kept in darkness for days before conducting spectroscopic experiments. Additionally, sample preparation was done in the dark, and the cuvette was covered whenever exposure to light was possible.

**Fig. 2.**
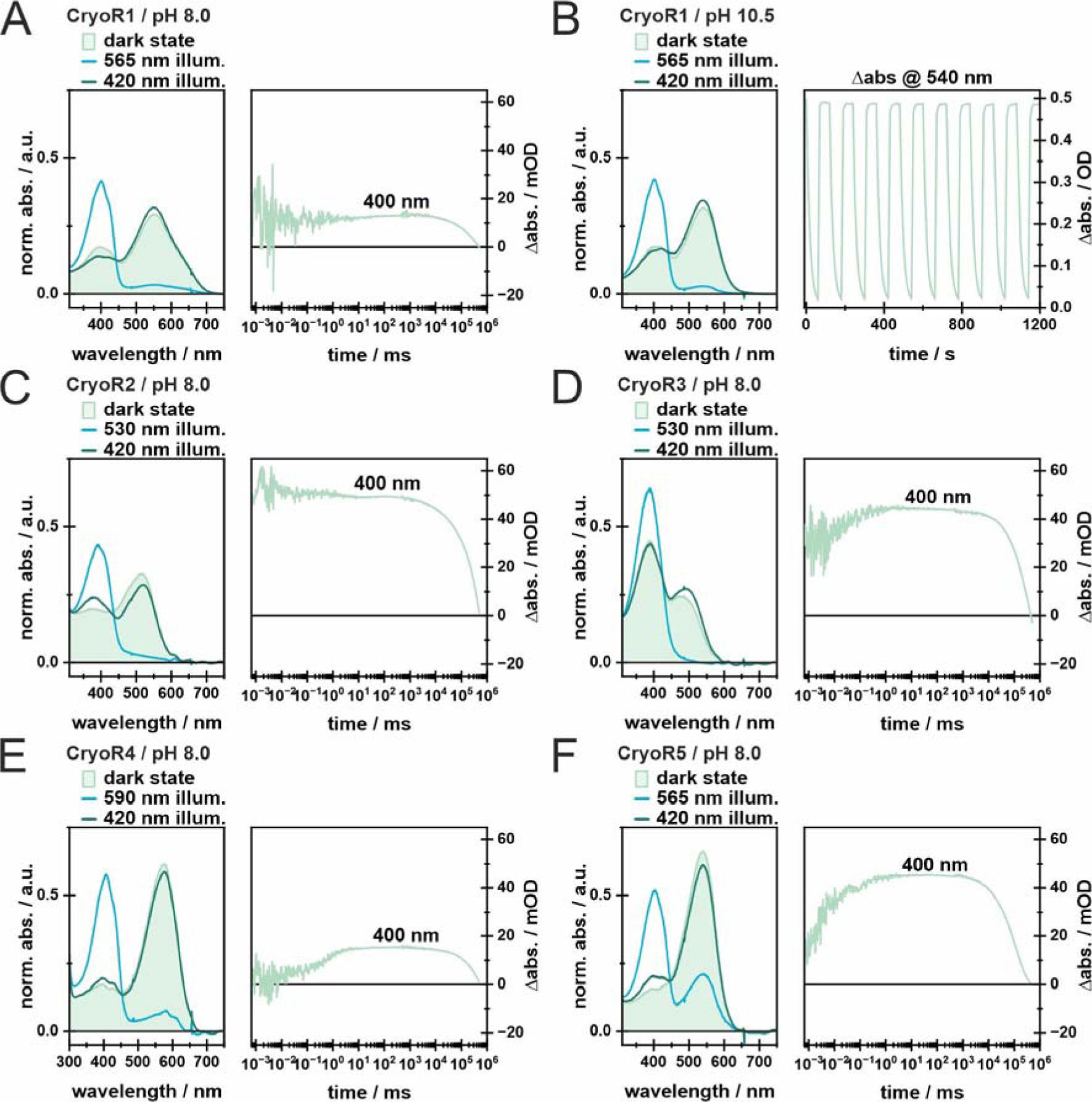
Illumination experiments and M state kinetics of CryoRs. Panels **A., C.-F.** show the dark state spectra (light green), as well as the PSS spectra after illumination of the main absorption band (blue) and the spectra after illumination of the obtained PSS with blue light (dark green) to recover the dark state of each investigated cryoR at pH 8.0. Each LED was turned on for 100 s. LEDs in the 500 nm range were operated at 100 mA, while the 420 nm LED was operated at 10 mA. Additionally, the transients at 400 nm are shown for each protein to elucidate the unusually slow decay of the M intermediate and the photocycle. Panel **B.** shows illumination experiments of CryoR1 at pH 10.5 including an alternating illumination scheme of green and blue-light to test for potential photobleaching and photofatigue. For all shown measurements, due to the very long photocycle duration and significant dark-state absorption at ∼400 nm, the very late signals may be come distorted due to actinic effects of the probe light.

For neutral to alkaline conditions illumination of all studied CryoRs with LEDs in the range of 530 nm to 590 nm, adjusted to the absorption maximum of the respective protein, resulted in a photostationary state (PSS) with an almost completely depopulated main absorption band and a newly formed UV signal centered at ∼400 nm (Fig. 2; Fig. S2B). At pH 8.0, for all CryoRs except CryoR3, the PSS peak shows a clearly-resolved fine structure with three distinguishable maxima. As shown for some other MRs, such a retro-retinyl-like absorption spectrum most likely originates from enforced planarity of the retinal chromophore^21–23^. Illumination of the PSS with a 420 nm LED resulted in the recovery of the dark state spectrum, which is well-known for MRs as the blue-light quenching effect (BLQ)^24–27^ (Fig. 2). Alternating illumination of the main absorption band and the populated PSS resulted in no photofatigue or photobleaching allowing the usage of BLQ to repopulate the parent state during experiments and potential applications in the future (Fig. 2B).

Without BLQ, CryoRs show unusually slow photocycle kinetics dominated by a blue-shifted intermediate, it takes several minutes to recover their parent state (Fig. 2). This is a characteristic feature of all studied CryoRs with various motifs. Thus, we propose it to be a common feature of the entire group.

In CryoR1, there are two such blue-shifted intermediates, M_1_ and M_2_, with lifetimes of ms and minutes, respectively (Fig. 3B; Fig. S3). The M_2_ state is slightly red-shifted compared to the M_1_ intermediate (Fig. 3B; Fig. S3). The M_1_ state is already populated at the beginning of the experimentally accessible flash photolysis timescale (Fig. 3B). Thus, the formation of the M_1_ state is accelerated compared to most MRs, where this transition happens on the µs to ms timescale^22,28,29^. Ultrafast spectroscopy data showed the formation of a red-shifted K intermediate preceding the M_1_ state (Fig. 3A; Fig. S4). Due to the experimental time gap on the ns timescale, we could not determine whether the M_1_ intermediate is directly populated after the K state, or if there is a multi-step K-to-M_1_ transition, as it was observed for various MRs previously, e.g. *Ns*XeR^21^(Fig. 3A,B). The resulting CryoR photocycle scheme is depicted in Fig. 3C, and is similar to the reported photocycle of DSE rhodopsins^14^.

**Fig. 3.**
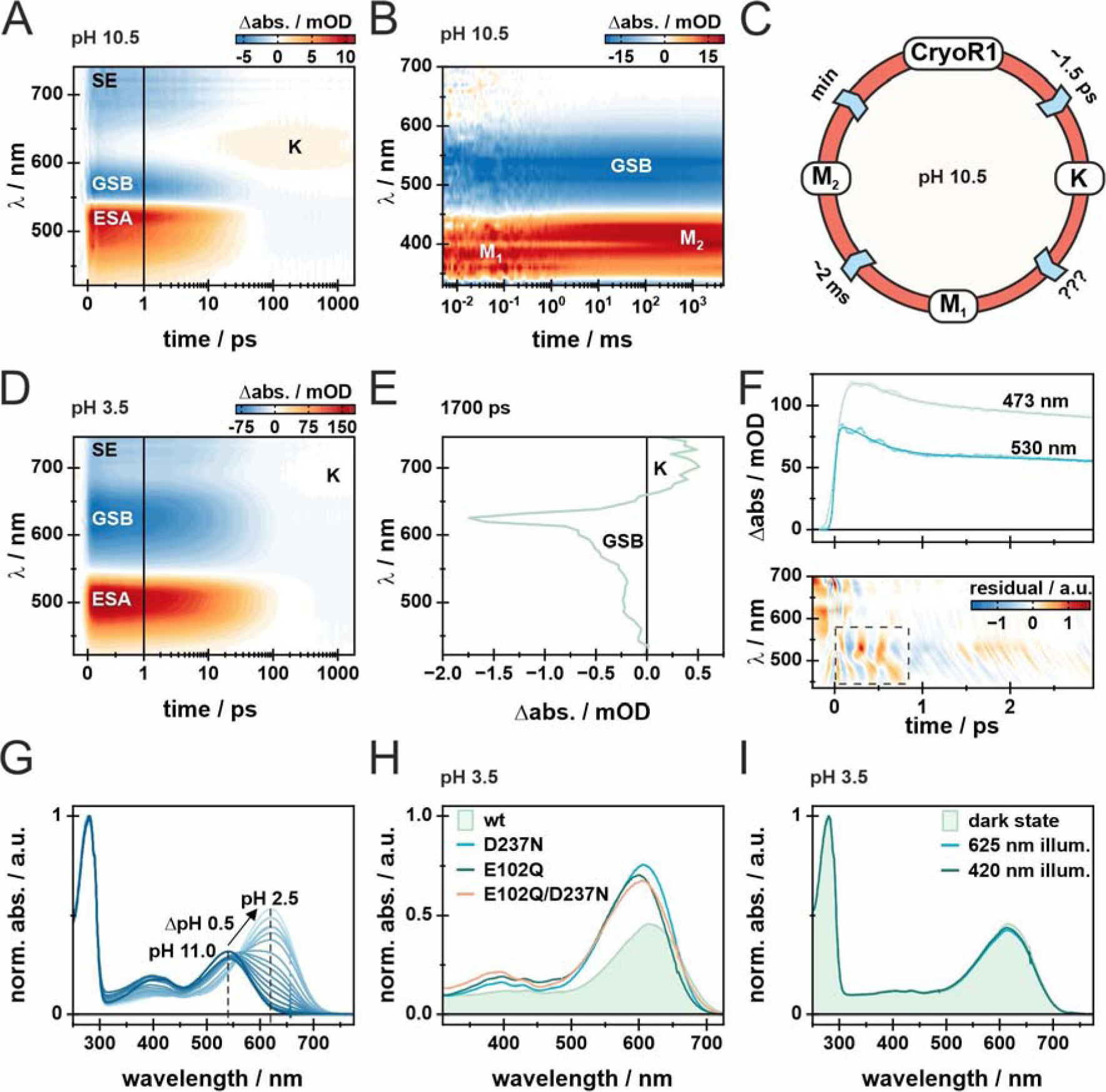
Spectroscopic characterization of CryoR1. **A.** Ultrafast transient absorption (TA) measurement of CryoR1 at pH 10.5. The signal amplitude is color coded as follows: positive (red), no (white) and negative (blue) difference absorbance. The x-axis is linear until 1 ps and logarithmic afterwards. **B**. Flash photolysis measurement of CryoR1 at pH 10.5 shown as a 2D-contour plot. The amplitudes are color coded in the same way as for the ultrafast measurement. The photocycle was measured until 5 s and quenched afterwards via illumination with a 405 nm LED (300 mA) before each acquisition. **C.** Photocycle model for CryoR1 at pH 10.5. **D.** Ultrafast TA measurement of CryoR1 at pH 3.5 shown as a 2D-contour plot. **E.** Difference spectrum after 1700 ps delay time taken from the fs TA measurement shown in panel **D**, showing the small amplitude for K formation and GSB once the photocycle is entered. The large negative amplitude at ∼620 nm is attributable to scattering of the excitation beam. **F.** Transients at two characteristic wavelengths (473 nm and 530 nm) taken from a fs TA measurement at pH 3.5 recorded and displayed on a linear timescale up to 3 ps and a decreased linear step size compared to the measurement shown in **D**. The raw data is displayed as dots and the obtained fit curves as lines to illustrate the observed coherent oscillations. Furthermore, the residuals of the raw experimental data and the fit are shown. The region of interest is highlighted with the dashed box. **G.** pH titration of the CryoR1 dark state absorption spectrum in the pH range 2.5 to 11.0. Spectra have been normalized with regard to the absorption band at 280 nm. **H.** Dark state absorption spectra of CryoR1 wt, D237N, E102Q, and E102Q/D237N mutants. **I.** Illumination experiments of CryoR1 performed at pH 3.5. The dark state was first illuminated with a 625 nm LED to check for a potential M state formation and with a 420 nm LED afterwards to check for potential BLQ-effect.

At acidic pH, the photocycle kinetics is accelerated for all studied CryoRs except CryoR3 (Fig. S5). Moreover, the spectral changes observed upon illumination were less pronounced compared to neutral and alkaline conditions (Fig. S5). Specifically, CryoR1, CryoR2, CryoR4, and CryoR5 show only minor changes in the absorption spectrum upon illumination of the main absorption band at acidic pH. The exception is CryoR3, still showing a formation of the blue-shifted PSS, but less intense compared to neutral and alkaline conditions (Fig. S5).

### Unique spectral behavior of CryoR1

Among the studied CryoRs, CryoR1 demonstrated several unique spectroscopic features, both in steady-state and time-resolved experiments. Therefore, we investigated the spectroscopic properties of CryoR1 in more detail.

The first feature is the ultrafast dynamics of CryoR1, being unusual at all studied pH values. At high pH, it resembles the pattern observed for various known MRs at low pH conditions or for their counterion mutants (Fig. 3A)^30–35^. Directly after excitation of the retinal chromophore, the excited-state absorption (ESA) (420-540 nm), ground-state bleach (GSB) (540-625 nm), and stimulated emission (SE) (625-740 nm) are observed. The ESA decays bi-exponentially (∼0.4 ps and ∼10 ps), with the faster lifetime showing the K intermediate formation in the range of ∼600-675 nm. In contrast to many known MRs, the typical J intermediate is missing.

At low pH, the ultrafast dynamics again show ESA, GSB, and SE signals but are dominated by a long-living ESA also decaying bi-exponentially (∼0.5 and ∼25 ps) as described by two-lifetime distributions in the corresponding lifetime distribution map (LDM) (Fig. 3D; Fig. S4A). However, the formation of the K intermediate is negligible (Fig. 3D,E). Interestingly, the wings of the ESA and GSB signals show strong coherent oscillations with two major frequency contributions, 32 cm^-1^ and 128 cm^-1^ and are damped on a ∼0.8 ps timescale (Fig. 3F; Fig. S6). The 32 cm^-1^ frequency corresponds to an oscillation period longer than the duration of the observed coherent oscillations. Therefore, this frequency is assigned to be an artifact of the Fourier analysis. The 128 cm^-1^ mode additionally shows contributions from slightly lower or higher frequencies (Fig. 3F; Fig. S6). Frequencies in this range have been linked to low-energy torsional modes of the C=C double bonds^36,37^. A phase shift of π is observed at the two wings of the ESA band, which fits well to a model of vibrational wave packet motion on the excited-state potential energy surface periodically modulating the ESA band (Fig. 3F). The strength of the coherent oscillations suggests a hindered retinal isomerization after excitation, in agreement with the extremely small signal amplitudes of the K state (Fig. 3D,E).

The second feature of CryoR1 is an unusually large spectral red shift of ∼80 nm upon pH titration in the range of pH 11.0 to pH 2.5 (Fig. 3G; Fig. S7). Specifically, at pH<6 the spectrum is dominated by the red-shifted band at 620 nm (Fig. 3G). To the best of our knowledge, this is the farthest red-absorbing non-eukaryotic MR reported to date. Additionally, under alkaline conditions, a second absorption band centered at 400 nm arises in the CryoR1 spectrum, which likely corresponds to the ground state species with a deprotonated RSB, due to its rise at low proton concentrations^38–40^. In contrast to many known MRs, the pH-dependent spectral shift of CryoR1 is not observed in a continuous movement across the spectral range^11,35,41^, but rather in a gradual decrease of the red-shifted absorption maximum at 620 nm associated with a rise of the blue-shifted absorption maximum at 540-550 nm and the deprotonated RSB band (400 nm). The large spectral shift of the observed maxima allows separated optical excitation of the respective species of interest.

Usually, pH-dependent shifts of the absorption maximum are associated with the change in protonation of the RSB or/and charged residues in the retinal binding pocket. Neutralization of such charged residues results in a red-shift of the absorption maximum, as it was observed for various MRs in the past^22,31,33^. Since in CryoR1 there are two RSB counterions, E102 and D237, we investigated the spectral properties of its E102Q, D237N, and E102Q/D237N variants (Fig. 3H; Fig. S8). None of the substitutions was sufficient to mimic the red-shift of the absorption maximum observed in the wild-type (WT) protein upon pH decrease (Fig. 3I). Indeed, the absorption maximum of the mutants was blue-shifted (10 - 15 nm) compared to that of WT at low pH. Therefore, the unique large spectral shift shown for CryoR1 cannot be simply described by the protonation of the RSB counterions and the molecular basis of this shift is further characterized below.

To gain insights into the architecture of CryoRs and their unique spectral properties we obtained high-resolution structures of CryoR1 and CryoR2 using single-particle cryo-EM and X-ray crystallography (Table S2; Supplementary Note 1). Both proteins form pentamers in detergent micelles, lipid nanodiscs, and crystals grown using the lipidic cubic phase (LCP) method (Fig. 4A; Supplementary Note 1). An elongated C terminus is capping the central part of the oligomer at the intracellular side (Fig. 4B,E). The C terminal elongation is conserved across CryoRs (Fig. S1). With this cap, the profile of the central pore of the CryoR pentamer is unusual compared to those of other known MRs (Fig. 4C,F). The transmembrane part has a narrow region near the W40 and R43 residues in CryoR1 (Y41 and R44 in CryoR2) (Fig. 4C,F), which to some extent resembles that of the pentameric viral rhodopsin OLPVRII^10^. However, the cytoplasmic part of the pore in CryoRs has an additional second narrowing formed by the C terminus near residues I238 and E287 in CryoR1 and CryoR2, respectively (Fig. 4B,C,E,F). Interestingly, the pore is completely blocked in this area in CryoR2 at acidic pH by five glutamic acid residues (E287) (Fig. 4F). The roles of such an unusual central channel and the unique cap formed by the C terminus remain elusive. Although the C terminus interacts with the nearby protomer within the oligomeric assembly and thus likely stabilizes the pentamer (Fig. S9), both the spectral properties of the truncated variant of CryoR1 (CryoR1_1-263_, 1-263 amino acid residues) including the long-living blue-shifted state as well as the oligomeric state are similar to those of the WT protein (Fig. 4G,H). Therefore, the presence of the elongated C terminus is likely not connected to the unique photocycle kinetics of CryoRs.

**Fig. 4.**
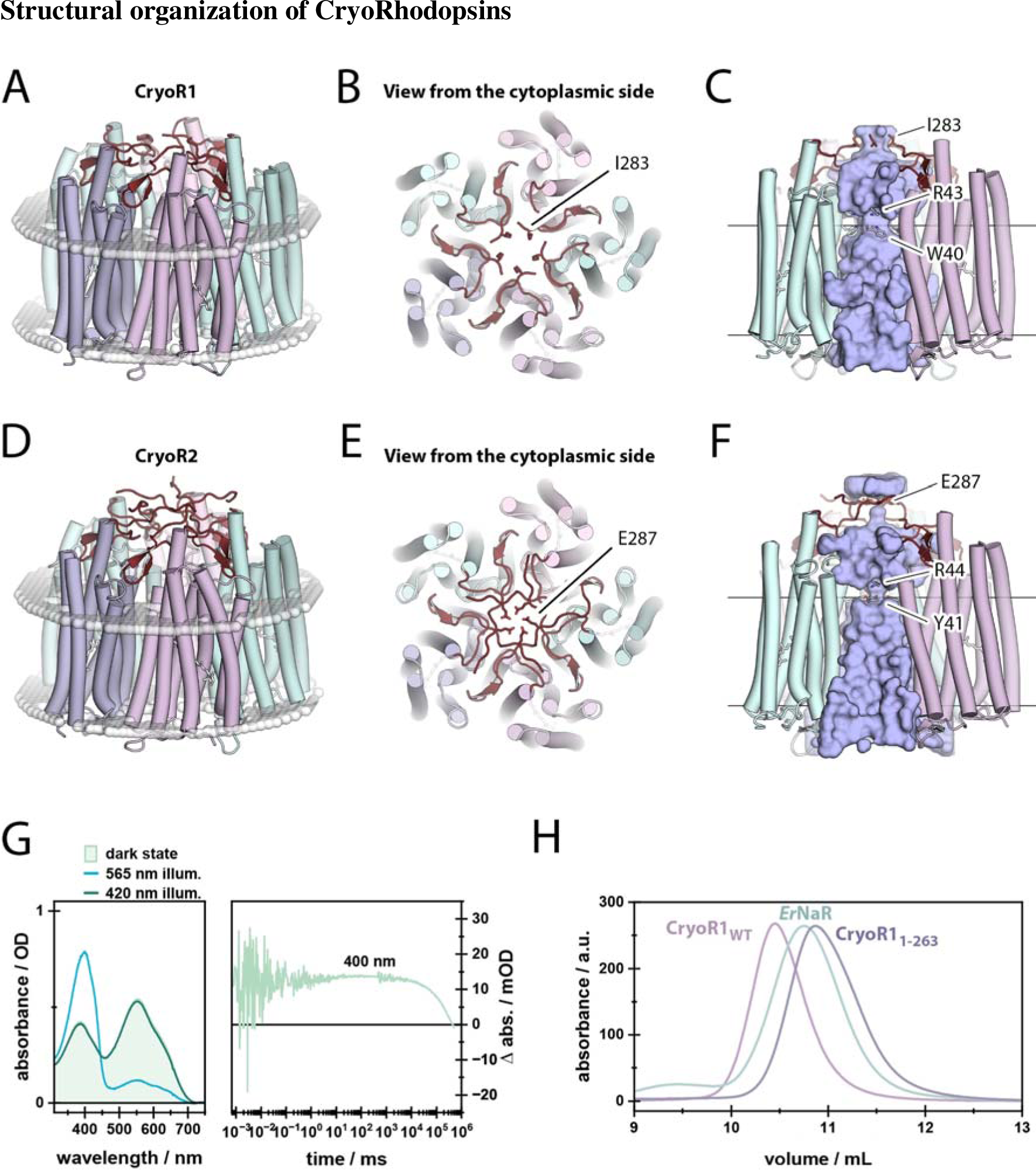
Pentameric architecture of CryoRs and unusual central channel. **A.** Overall view of the CryoR1 pentamer. **B.** View at the CryoR1 pentamer from the cytoplasmic side. **C.** Side view of the central channel in CryoR1. **D.** Overall view of the CryoR2 pentamer in crystals. **E.** View at the CryoR1 pentamer from the cytoplasmic side. **F.** Side view of the central channel in CryoR1. C terminus is colored dark red. **G.** Spectroscopy of CryoR1_1-263_. Left: the dark state spectra (light green), as well as the PSS spectra after illumination of the main absorption band (blue), the spectra after illumination of the obtained PSS with blue light (dark green) to recover the dark state of CryoR1_1-263_ at pH 8.0. The 565 nm LED was turned on for 100 s at 100 mA, while the 420 nm LED turned on for 100 s at 10 mA. Right: the transient at 400 nm. **H.** Size-exclusion chromatography (SEC) profiles of pentameric CryoR1_WT_ (MW: 180.6 kDa) and CryoR1_1-263_ (MW: 147.2 kDa). SEC profile of the pentameric light-driven sodium pump *Er*NaR (MW: 164.0 kDa) measured under the same conditions using the same column is shown as a reference.

Another interesting feature of CryoRs is a cavity at the cytoplasmic side found in each interprotomeric cleft and also restricted by the β-sheet of the C terminus and the hydrophobic/hydrophilic boundary of the intracellular membrane leaflet (Fig. 5A,B). The cavity goes through a pore between helices A and G and leads to another cavity inside the protomer (Fig. 5B). The latter cytoplasmic cavity is filled with several water molecules and is surrounded by positively charged conserved amino acid residues, R38, H249, and R57 (CryoR1 numbering), which is the characteristic arginine of CryoRs (Fig. 5B,C). In the ground state of CryoR1, R57 is shielding the E113 residue of the functional motif from the cytoplasmic cavity (Fig. 5B,C).

**Fig. 5.**
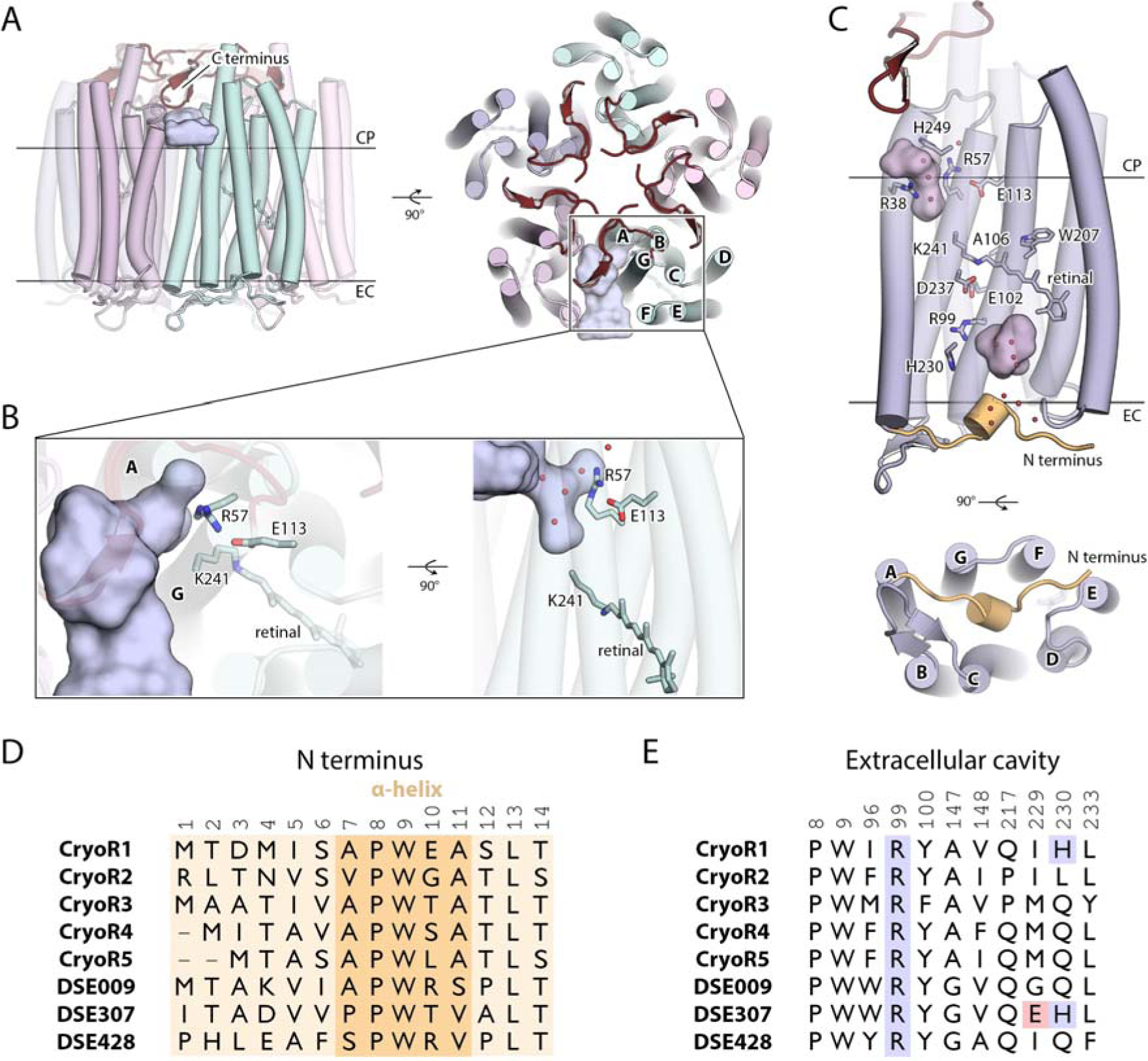
Structural features of the CryoR protomer. **A.** Side view (left) and view from the cytoplasmic side (right) of the CryoR1 pentamer and the interprotomeric cavity (shown with light blue surface). **B.** Detailed view of the interprotomeric cavity and the internal cytoplasmic cavity in CryoR1. **C.** Side view (top) and view from the extracellular side (bottom) of the protomer of CryoR1. N terminus is colored wheat. The small α-helix in the N terminus is shown as a cylinder. Internal cavities are colored light pink. **D.** Sequence alignment of the N terminus region of representative CryoRs and DSE rhodopsins. **E.** Alignment of the amino acids surrounding the extracellular cavity of representative CryoRs and DSE rhodopsins.

In contrast to all known MRs, there are no internal water molecules in the central region of CryoR1 and CryoR2 (Fig. 5C; Supplementary Note 1). Indeed, no waters were observed even at the cytoplasmic side of the retinal binding pocket, where normally at least one is found in other MRs near the highly conserved tryptophan residue of the helix F (W207/W208 in CryoR1/CryoR2, W182 in BR^42,43^). In the RSB region of both CryoR1 and CryoR2 up to the arginine residue at the extracellular side (R99/R100 in CryoR1/CryoR2) there are also no internal water molecules. In total, no water molecules were found in the radius of ∼11 Å from the RSB, which is a unique feature of the studied CryoRs. The absence of the water molecules in the RSB region was also shown for rhodopsin phosphodiesterase (Rh-PDE) from *Salpingoeca rosetta*^44^. The set of counterions of Rh-PDE is identical to CryoR1 and CryoR2 (E164 and D292 in Rh-PDE). However, in contrast to CryoRs, in Rh-PDE there is a water molecule at the cytoplasmic side of the retinal binding pocket similar to other MRs. Thus, we suggest that the absence of internal water molecules in the central part of the protein molecule might be a feature of CryoRs.

At the extracellular side of CryoRs, we found a large water-filled cavity, between R99/R100 (in CryoR1/CryoR2; analog of R82 in BR) and the small (single turn) N terminal α-helix, capping the internal space of the CryoR protomer from the extracellular bulk (Fig. 5C; Supplementary Note 1). This helix is similar to that of the light-driven sodium pumps KR2^45,46^ and *Er*NaR^47^ and likely other NDQ rhodopsins but in CryoRs it is shorter. Both the extracellular cavity and the N terminal α-helix are likely conserved within CryoRs and are found as well in DSE rhodopsins (Fig. 5D,E). Aside from the R99/R100, there are no internal rechargeable residues at the extracellular side of CryoRs. The exception is H230 in CryoR1 (Fig. 5C), which is only found in 2 out of 40 members of the CryoRs group. The presence of the water-filled cavity may provide a weak transport of ions through the region in accordance with the slow photocycle kinetics in a similar way to that recently proposed for an inward proton pump *Bc*XeR^48^.

### Newly identified arginine flips in the blue-shifted state

As it can be concluded from the spectroscopic analysis, CryoR1 can be photoconverted at high pH between the ground and the M_2_ states (Fig. 6A). Accordingly, we prepared the CryoR1 sample at pH 10.5 in both the ground and the M_2_ states using pre-illumination with 405 nm and 530 nm LED lamps, respectively, for the cryo-EM studies (Fig. 6A). The pre-illuminated samples were applied to the cryo-EM grids and vitrified in liquid ethane. In the case of the ground state of CryoR1, to preserve the purity of the state, the grids were prepared under 405 nm LED lamp illumination (Fig. 6B). The resulting cryo-EM maps at 2.7 Å resolution showed the presence of a single state, which we assigned to the ground-state (Fig. 6C,D).

**Fig. 6.**
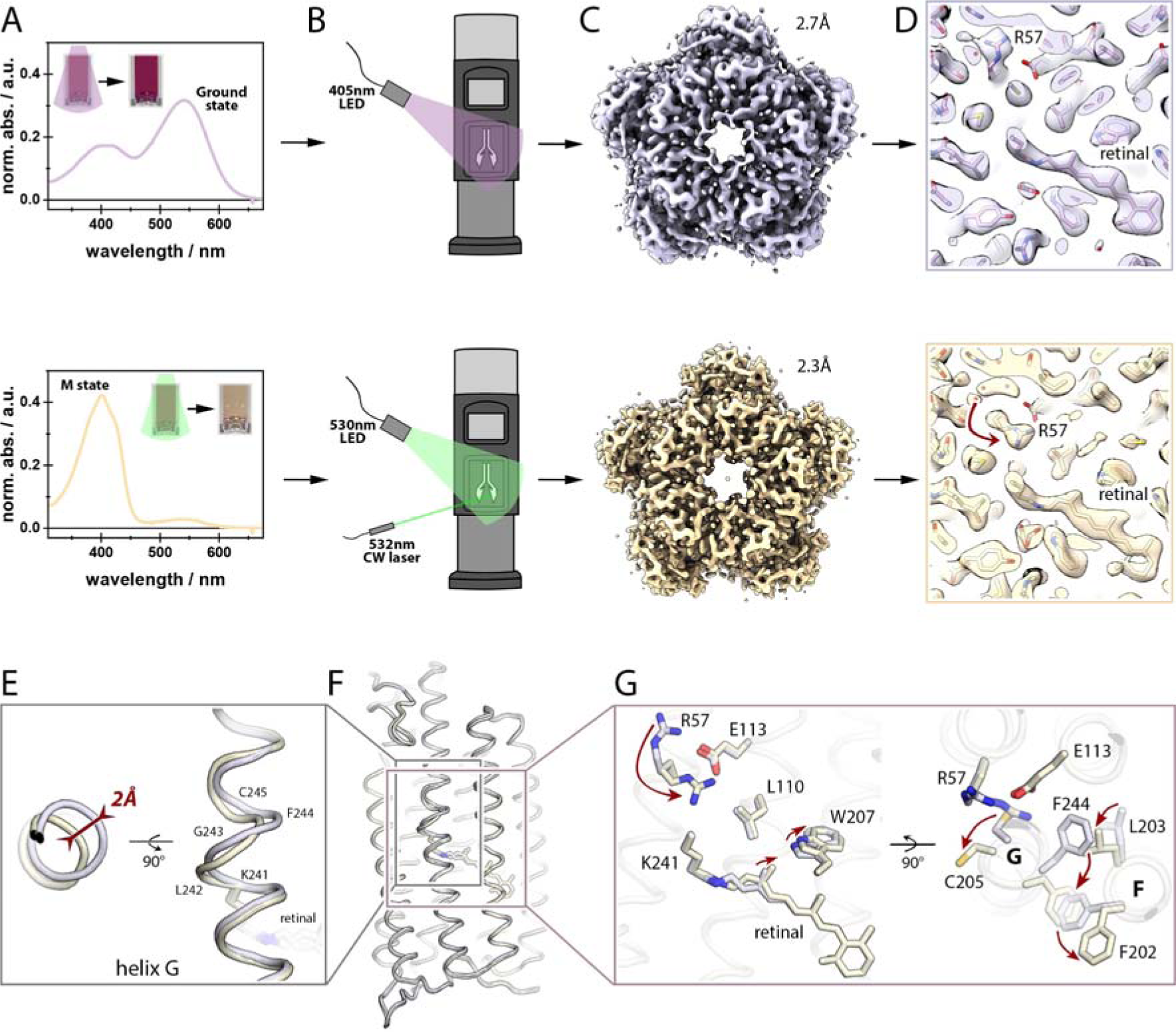
Cryo-EM structures of the ground and M_2_ states of CryoR1. **A.** Preillumination of the sample with the 405 and 530 nm LEDs for the ground (top) and M_2_ (bottom) states, respectively. The insets show the sample before and after illumination. Spectra correspond to the illuminated samples. **B.** Scheme of the cryo-EM grid preparation. For the ground state, the grid was prepared under the 405-nm LED light (top). For the M_2_ state, the grid was prepared under the 530 nm LED and 532nm laser light (bottom). **C.** Cryo-EM maps of the ground (top) and M_2_ (bottom) states of CryoR1. **D.** Cryo-EM maps near the retinal and R57 in the ground (top) and M_2_ (bottom) states. The R57 rearrangement is indicated with a red arrow. **E.** Distortion of helix G in the M_2_ state (light yellow) compared to the ground state (light purple). Maximum displacement is indicated with a red arrow. **F.** Overall alignment of the CryoR1 protomer in the ground (light purple) and M_2_ (light yellow) states. **G.** Detailed view of the rearrangements of R57 and residues in helices F and G between the ground (light purple) and M_2_ (light yellow) states. The major changes are indicated with red arrows.

In the case of the accumulated M_2_ state, to maximize the occupancy of the intermediate, the grids were prepared under green light illumination using both a 530 nm LED and a 532 nm laser beam centered onto the grid through the sample application hole on the side of the vitrification robot (Fig. 6B). The resulting cryo-EM maps at 2.3 Å resolution revealed the presence of at least two states of the protein (Fig. 6C,D). The first is minor and similar to the ground state, while the second is more dominant and clearly different; we assigned it to the M_2_ state.

The cryo-EM map near the retinal and the RSB strongly suggested the 13-*cis* conformation of the cofactor in the M_2_ state (Fig. 6D-G). The side chain of W207 is rotated compared to the ground state to allow space for a shifted by 0.6 Å C20 atom of the retinal, also indicating the isomerization of the cofactor (Fig. 6G). Following the retinal isomerization, the central part of helix G is distorted in the M_2_ state in the region of residues K241-C245 (Fig. 6E,F). It is accompanied with significant flips of the side chains of F244 and C245 (Fig. 6G). The large-scale movement of F244 occurs synchronously with the reorientations of F202 and L203 in the helix F (Fig. 6G).

The above-mentioned structural rearrangements at the cytoplasmic side of CryoR1 are associated with the flip of the unique characteristic R57 residue towards the RSB (Fig. 6G). While in the ground state R57 points to the cytoplasmic side, in the M_2_ state it is oriented towards the center of the protomer and is located within ∼6.5 Å of the RSB. Notably, in both the ground and M_2_ states R57 forms a salt bridge with E113 of the functional motif and is additionally stabilized by the main chain of helix G. E113 remains at the same position in the ground and M_2_ states (Fig. 6G).

The X-ray crystallography and cryo-EM data on CryoR2 also showed the flip of the characteristic arginine accompanied by the similar rearrangements of the residues in helices F and G (Supplementary Note 1; Fig. S11, S12). In that case, R58 (analog of R57 in CryoR1) forms a salt bridge with E114 (analog of E113 in CryoR1) in the ground and M_2_ states of CryoR2 in almost identical manner to those shown for CryoR1. Thus, structural data on CryoR1 and CryoR2 allowed us to conclude that the reorientation of the characteristic unique arginine at the cytoplasmic side of CryoRs in the long-living blue-shifted state is likely a feature of the entire group.

### Wavelength- and voltage-dependent proton translocation in CryoR1

Ion-transporting activity tests of the selected proteins by measuring pH changes in the *E.coli* cell suspension upon illumination did not report ion transport (Fig. S10). The results remained the same in the presence of protonophore carbonyl cyanide m-chlorophenyl hydrazone (CCCP), indicating the absence of detectable transport activity at least for protons, chloride, and sodium ions (Fig. S10). The results of ion-transport assays are in line with the extremely slow photocycle kinetics, not allowing the detectable pH changes in the cell suspension.

To get insights into the proton transport mechanism of CryoRs, we next measured photocurrents in NG108-15 expressing CryoR1 in whole-cell patch-clamp experiments. Since CryoR1 has a broad absorption band at pH 7.4, we first used the 565 nm nanosecond light pulses to activate the co-existing subpopulations of the rhodopsin. We obtained transient photocurrents, the vectoriality of which changed depending on the applied voltage (Fig. S11A).

However, we were unable to elicit stationary photocurrents without additional activation of the long-living M_2_ state, due to the very slow RSB reprotonation kinetics (minutes). The activation of the M_2_ state accelerates the following transition to the ground state (Fig. 2). Hence, we employed a white light source, which also included a blue light component, allowing for simultaneous activation of the ground and the M_2_ states^24–27^. As a result, we obtained negative stationary currents during white light illumination at voltages ranging from -80 to 40 mV (Fig. S11B), thereby demonstrating that CryoR1 exhibits an inward proton-pumping activity.

The spectroscopy of CryoR1 has revealed two ground state subpopulations at pH 7.4 with distinct absorption maxima, one at 550 (likely for the non-protonated RSB counterion complex) and 620 nm (protonated RSB counterion complex) (Supplementary Note 2). Thus, to further investigate the mechanism of inward proton pumping and vectoriality change in CryoR1, we separately activated these two ground state subpopulations and the M_2_ state by employing nanosecond pulses of 500, 620, or 420 nm, respectively.

The activation of the ground state subpopulation with the deprotonated counterion complex using 500 nm pulses led to positive transients (Fig. S11C). The positive transient currents reflect proton transfer towards the extracellular side, likely from the RSB to the counterion complex (Fig. S12A).

On the contrary, the activation of the ground state subpopulation with the protonated counterion complex using 620 nm pulses resulted in negative transients at all voltages (Fig. S11D). Such negative transient currents indicate proton transfer to the cytoplasmic side. Thus, when the counterion complex of CryoR1 is protonated, the RSB proton is likely transferred to either E113 or directly the cytoplasmic bulk through the internal cavity (Fig. S12B).

The activation of the M_2_ state with 420 nm light pulses resulted in negative transient photocurrents at voltages ranging from -40 to +40 mV (Fig. S11E). We integrated the transient photocurrents to quantify the absolute total charge transfer in the transition from M_2_ to the ground state, and in the transitions from the two ground state subpopulations to the M states normalized by their respective population fractions. This analysis revealed that the M_2_-to-ground state transition involves the largest amount of charge transfer (Fig. S13), likely through the reprotonation of the RSB, the subsequent partial reprotonation of the counterion complex, as well as the flip of R57 towards the cytoplasmic side observed in the structural data. Notably, as the voltage was changed from -40 to +40 mV the absolute values of the transient amplitudes increased for 500 nm activation and decreased for 620 nm activation. This together with the photocurrent reversal at nearly 0 mV upon 565 nm activation suggests that the distribution between the two ground state subpopulations is voltage-dependent. A similar proton transport mechanism was previously described for a proteorhodopsin (PR), which showed photocurrent reversal at acidic conditions (pH 5.5) at a negative membrane potential of -80 mV^49^.

### Determinants of the dominant blue-shifted state of CryoRs

Our data suggest that the dominant blue-shifted state is a key element in CryoRs, and maximizing its population could be vital for the proper functioning of CryoRs under harsh conditions. To make the state dominant, two features have to be fulfilled: (*1*) this intermediate should be effectively formed and (*2*) it should be stable enough to decay very slowly. The strong blue shift of the spectra in the M_1_ and M_2_ states reflects the deprotonated form of the RSB. Therefore, to satisfy both conditions, the RSB in CryoRs should be effectively deprotonated and very ineffectively reprotonated. Below we describe the mechanisms that were evolutionarily developed in CryoRs to naturally maximize the lifetime of the blue-shifted states.

The deprotonation of the RSB proceeds quickly in all studied CryoRs (Fig. 2). Our structural data strongly suggest that the proton from the RSB is transferred to the complex of two counterions (E102-D237 in CryoR1), which directly interact in the M_2_ state at pH 10.5 (Fig. 7A,B; Fig. S16; Supplementary Note 2). It is well-known that such pairs of carboxylic residues may have a very high proton affinity allowing them to effectively store the proton released from the RSB and to hamper its backflow.

**Fig. 7.**
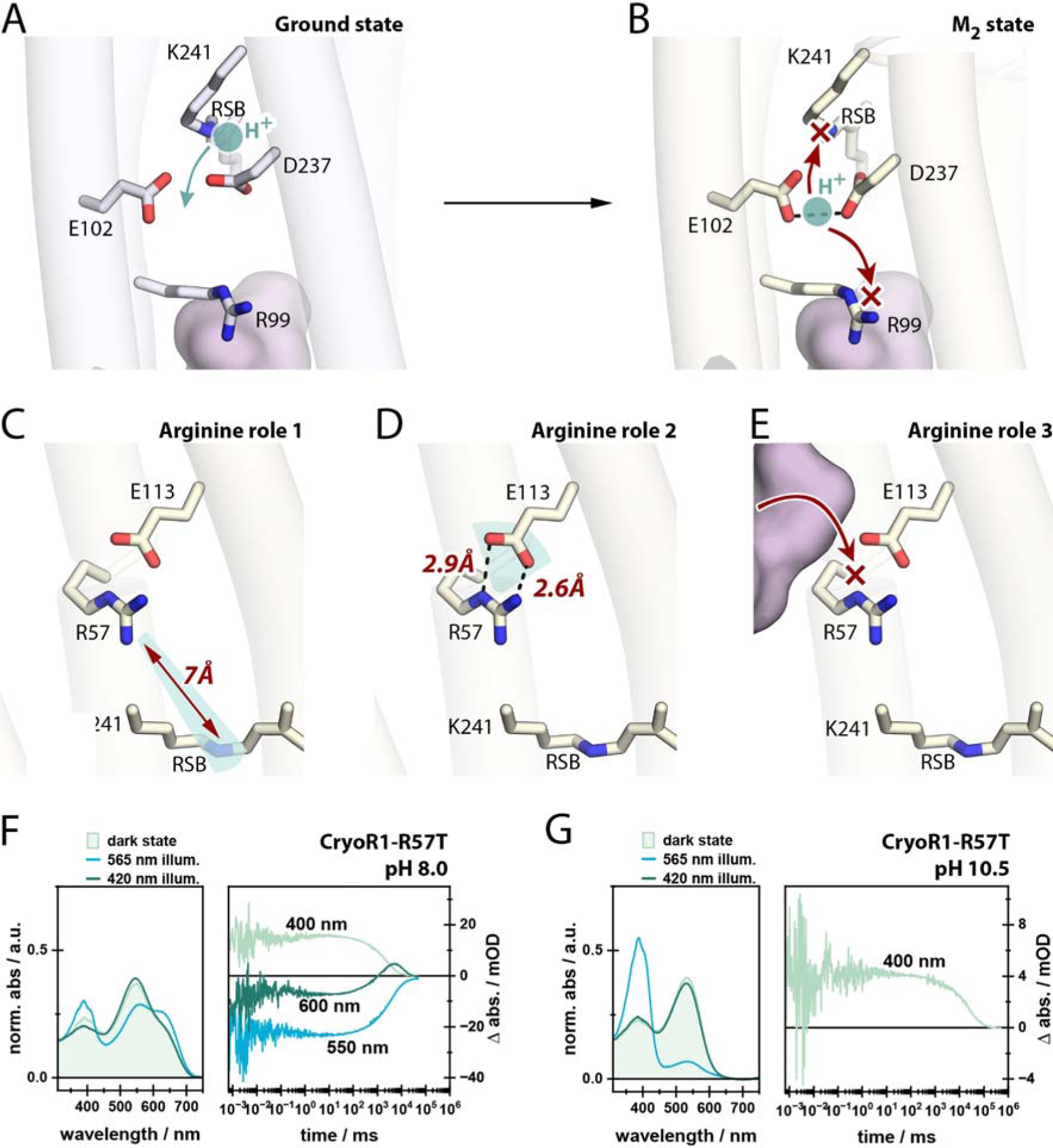
Determinants of the dominant M state in the CryoR photocycle. **A.** The RSB region in the ground state of CryoR1 at pH 10.5. The proton at the RSB is indicated with a cyan circle. The tentative proton relocation with the rise of the M state is indicated with a cyan arrow. **B.** The RSB region in the M state of CryoR1 at pH 10.5. The proton relocated to the E102-D237 complex is shown with a cyan circle. The blocked transfer pathways of the proton to the RSB and towards the extracellular side are indicated with red arrows and crosses. **C.** R57 effect on the pKa of the RSB in the M state of CryoR1 (region of the effect highlighted cyan). **D.** R57 effect on the pKa of E113 in the M state of CryoR1 (region of the effect highlighted cyan). **E.** R57 blocks the pathway for protons from the cytoplasm to the RSB. The cytoplasmic cavity is shown with a pink surface. **F.** Spectroscopy of CryoR1_R57T_ at pH 8.0. Left: the dark state spectrum (light green), as well as the PSS spectrum after illumination of the main absorption band (blue) and the spectrum after illumination of the obtained PSS with blue light (dark green) to recover the dark state. Right: the transients at 400, 550, and 600 nm. **G.** Spectroscopy of CryoR1_R57T_ at pH 10.5. Left: the dark state spectrum (light green), as well as the PSS spectrum after illumination of the main absorption band (blue) and the spectrum after illumination of the obtained PSS with blue light (dark green) to recover the dark state. Right: the flash photolysis transient at 400 nm. The 565 nm LED was turned on for 100 s at 100 mA, while the 420 nm LED was turned on for 100 s at 10 mA.

At the same moment, the RSB reprotonation is greatly slowed in CryoRs. Based on our experimental data it can be explained as follows. One reason originates from the above-mentioned proton acceptor complex (E102-D237 in CryoR1) and the architecture of the extracellular side of the group members (Supplementary Note 2). Indeed, the proton released from the RSB to its counterions can barely be transferred further as there is an arginine (R99 in CryoR1) barrier and no rechargeable internal groups at the extracellular side of CryoRs (Fig. 5C,E; Fig. 7B). The region is also isolated from the bulk by the cap formed by the N terminal α-helix (Fig. 5C). Thus, the proton is likely trapped at the counterion complex in the M_2_ state (Fig. 7B). At the same time, our spectroscopy studies of the E102Q, D237N, and E102Q/D237N mutants of CryoR1 as well as the E102Q mutant of CryoR2, imitating various protonation states of the counterion complex, clearly showed that once the complex gets protonated, the RSB deprotonates as evidenced by the strong blue shift of the absorption spectra (Fig. S8, Supplementary Note 2). Therefore, we suggest that the proton, released from the RSB and trapped at the counterion complex in the M_2_ state, stabilizes this long-living intermediate and can only be returned to the Schiff base after slow retinal reisomerization to the original conformation (Fig. 7B).

Another reason for the unusually slow RSB reprotonation is associated with an elegant use of the newly identified unique arginine residue at the cytoplasmic side (Fig. 7C-E). Indeed, our structural data show that in both CryoR1 and CryoR2, the arginine likely plays a triple role. First, by relocating towards the RSB in the M_2_ state, it lowers the pKa of the Schiff base, hampering its reprotonation even at neutral and mildly acidic pH values (Fig. 7C). Second, the arginine decreases the pKa of a conserved glutamic acid at the cytoplasmic side (E113/E114 in CryoR1/CryoR2) (Fig. 7D). For instance, in PRs the reprotonation of the RSB proceeds from the internal proton donor^50–52^, which is typically a glutamic acid residue at the cytoplasmic side (E108 in green PR, or GPR). This allows an effective RSB reprotonation at neutral and mild-alkaline pH values resulting in outward proton-pumping activity of PRs. In CryoRs, the analogous glutamic acid remains deprotonated at pH as low as 4.6 as indicated by the salt bridge formed between R58 and E114 in the crystal structure of CryoR2 (Fig. S17; Supplementary Note 1). Only at pH 4.3 in CryoR1, the bridge is broken, tentatively indicating the protonation of the glutamic acid (Fig. S17). The spectroscopy analysis of the E113Q variant of CryoR1 also showed no effect of the mutation on the photocycle kinetics (Fig. S18), strongly suggesting that the glutamic acid does not serve as a proton donor in CryoRs. Lastly, the arginine side chain in the M_2_ state blocks the pathway for a proton to the RSB from the cytoplasmic side serving as a positively charged barrier similar to that at the extracellular side (Fig. 7B,E). We suggest the latter role be the key for the observed inward proton-pumping modality of CryoR1, as R57 might prevent the backflow of the proton after its transfer from the RSB towards the cytoplasmic side.

To probe the roles of this arginine in the stabilization of the M_2_ state in CryoRs, we studied the spectroscopic properties of the R57T mutant of CryoR1. In agreement with the proposed role 1, the maximum absorption wavelength of CryoR-R57T differed from that of CryoR1-WT both at pH 8.0 (545 nm vs. 550 nm) and 10.5 (534 nm vs. 541 nm) already in the ground state (Fig. 7F,G; Fig. S19). Furthermore, the difference in the UV peak position between the mutant and WT is more pronounced in the PSS state (389 nm vs. 400 nm at pH 8.0 and 388 nm vs. 400 nm at pH 10.5), in line with a shorter distance (6.5 Å) between R57 and the RSB in the intermediate (Fig. 7C; Fig. S19). Next, at pH 8.0, the decay of the blue-shifted state in the mutant is ∼100 times faster than in the CryoR1-WT (Fig. 7F; Fig. S19). This coincides with the pronounced formation of the red-shifted late intermediate (Fig. 7F). At pH 10.5, the behavior is similar to that of WT (Fig. 7G). Thus, the data strongly suggests that the unique arginine of CryoRs stabilizes the blue-shifted state providing for its unusually long lifetime at neutral and mild-alkaline pH. At alkaline pH, the photocycle kinetics is also extremely slow but seems independent of the arginine. Such behavior is expected since the concentration of protons is much lower, which typically for MRs results in a notably decelerated photocycle.

## DISCUSSION

Here we characterized a newly identified group of MRs, CryoRhodopsins, members of which were found in cold-living bacteria. CryoRhodopsins demonstrate common features such as a minutes-long photocycle dominated by a blue-shifted intermediate state and the presence of the arginine residue at the cytoplasmic side near the position of the proton donor in many archaeal, bacterial, and eukaryotic proton-pumping rhodopsins.

Our data suggests that CryoRs exhibit inward proton-pumping activity. Notably, this requires illumination by white light sources containing a UV component to accelerate Schiff base reprotonation as exemplarily shown for CryoR1. Due to the requirement of the UV component for substantial proton pumping activity, we consider that CryoRs might play a role in sensing potentially harmful UV light.

The characteristic arginine in the helix B is the key determinant allowing CryoRs to dramatically increase the lifetime of the blue-shifted state. The molecular mechanism of the stabilization of the blue-shifted state involves the unusual flipping motion of the arginine residue to serve as a barrier for proton translocation back to the retinal Schiff base. Beyond this residue, the overall structure of CryoRs and unusual sets of counterions also contribute to stabilize this state.

The intriguing ability of retinal proteins to form a photochromic pair – comprising two forms with distinct absorption maxima and reversible conversion through photonic stimuli – has captivated scientists since early times. This fascination stems from the potential application of these proteins in recording, processing, and storing optical information, giving rise to numerous proposed uses such as three-dimensional optical memories, real-time holographic processors, and artificial retinas. While BR has historically been the most employed protein for these purposes, its principal drawback lies in the short lifetime of the blue-shifted M state. Various strategies^53,54^ have been employed to address this limitation, including chemical modification of retinal, site-specific mutagenesis, chemical supplements, directed evolution, or the utilization of branched P/Q-states – each accompanied by its unique drawbacks. In this context, CryoRs, boasting a naturally prolonged M state and heightened light sensitivity, presents itself as a promising template for future advancements in this field.

The ability of CryoR1 to elicit opposing transient photocurrents in response to ground state activation with red (620 nm) and green (500 nm) light pulses suggests potential utility of CryoRs for bidirectional, dual-color optogenetic control of cells, which currently involves the use of tandem protein constructs^55,56^ However, the partial charge transfers in CryoR1 upon illumination with either red or green light pulses are likely too small for optogenetic activation or silencing of excitable cells. Nevertheless, CryoR variants, which exhibit larger partial charge transfer, faster RSB reprotonation and strong plasma membrane targeted expression might allow bidirectional optogenetic control with only one oligomeric rhodopsin in the future.

Finally, the variation of functional motifs within the CryoR group, in particular, the alteration of amino acid residues close to the retinal cofactor, results in notably different absorption spectra of the studied proteins. Since the environments of the source organisms containing CryoRs vary significantly (glaciers, high mountains, boreal waters), we hypothesize that such an evolutional adjustment of the motifs is necessary for efficient functioning under these different light conditions. Taking into account the extreme and harsh environmental conditions of cold-living organisms like *Cryobacteria* and difficulty of their cultivation, our work provides a solid basis for further investigations of CryoRhodopsins and also sheds light onto the unusual light-sensing in psychrotrophic bacteria.

## METHODS

### Search for and phylogenetic analysis of CryoRhodopsins

Initial search for the novel rhodopsins was performed against the UniProtKB (2023-09-15), UniParc (2023-09-15), Genbank (2023-09-15), and MGnify (2023-09-15) databases using hmmsearch with the HMM profile of the microbial rhodopsins family from Pfam (PF01036). The 7-letter motifs were retrieved using custom Python scripts using jupyter notebook. Then, the final search was performed using jackhmmer (N iterations = 5) using the sequence of CryoR1 rhodopsin (GenBank ID: WP_166787544) as a template. The sequences possessing the RXXXXXK motif were selected, filtered to contain more than 250 amino acid residues, and re-aligned separately using MUSCLE^57^. Duplicated sequences were removed using a custom Python script. The selected sequences were also manually inspected to contain full seven transmembrane α-helices. The maximum likelihood phylogenetic tree was built using IQ-TREE webserver^58^ (Number of bootstrap alignments: 1000) and visualized using iTOL (v6.8)^59^. For the tree building we used a multiple sequence alignment (MSA) file containing 2199 sequences of microbial rhodopsins from ^1^. The sequences of CryoRhodopsins and three sequences of the DSE rhodopsins reported in ^14^ were added to the MSA file using MAFFT^60^. For the building of the phylogenetic tree all sequences with identity higher than 90% were removed using CD-HIT software^61^. The full MSA file, 90% identity-filtered MSA file, and phylogenetic tree file are provided as Supplementary files.

### Cloning

The DNA coding CryoR1-5 were optimized for *E.coli* using GeneArt (Thermo Fisher Scientific). Genes were synthesized commercially (Eurofins). For protein expression and purification, the pET15b plasmid with 6xHis-tag at the C-terminal was used. Point mutations and C terminus deletions were introduced using whole-plasmid PCR followed by the blunt ends ligation.

For electrophysiological recordings a human codon optimized gene of CryoR1 was cloned into the pcDNA3.1(-) vector between *BamH*I and *Hind*III sites together with an N-terminal part of channelrhodopsin (C2C1), membrane trafficking signal (TS) and ER export signal (ES) from potassium channel Kir2.1 and enhanced yellow fluorescent protein (EYFP). C2C1 and TS-EYFP-ES were amplified from pEYFP-N1-eKR2^62^, which was a gift from Prof. Peter Hegemann (Addgene plasmid #115337).

### Protein expression, solubilization, and purification

*E.coli* cells were transformed with pET15b plasmid containing the gene of interest. Transformed cells were grown at 37°C in shaking baffled flasks in an autoinducing medium ZYP-5052^63^, containing 10 mg/L ampicillin. They were induced at an OD_600_ of 0.8-0.9 with 1 mM isopropyl-β-D-thiogalactopyranoside (IPTG). Subsequently, 10 μM all-*trans*-retinal was added. Incubation continued for 3 hours. The cells were collected by centrifugation at 4000*g* for 25 min. Collected cells were disrupted in an M-110P Lab Homogenizer (Microfluidics) at 25,000 p.s.i. in a buffer containing 20 mM Tris-HCl, pH 8.0, 5% glycerol, 0.5% Triton X-100 (Sigma-Aldrich) and 50 mg/L DNase I (Sigma-Aldrich). The membrane fraction of the cell lysate was isolated by ultracentrifugation at 125,000*g* for 1 h at 4°C. The pellet was resuspended in a buffer containing 20 mM Tris-HCl, pH 8.0, 0.2 M NaCl and 1% DDM (Anatrace, Affymetrix) and stirred overnight for solubilization. The insoluble fraction was removed by ultracentrifugation at 125,000*g* for 1h at 4 °C. The supernatant was loaded on a Ni-NTA column (Qiagen), and the protein was eluted in a buffer containing 20 mM Tris-HCl, pH 8.0, 0.2 M NaCl, 0.4 M imidazole, and 0.1% DDM. The eluate was subjected to size-exclusion chromatography on a Superdex 200i 300/10 (GE Healthcare Life Sciences) in a buffer containing 20 mM Tris-HCl, pH 8.0, 100 mM NaCl and 0.03% DDM. In the end, protein was concentrated to 70 mg/ml for crystallization and –80°C storage.

### Measurements of pump activity in the *E. coli* cells suspension

CryoRs were expressed as described above. The cells were collected by centrifugation at 4,000*g* for 15 min and were washed three times with an unbuffered 100 mM NaCl solution, with 30-min intervals between the washes to allow for exchange of the ions inside the cells with the bulk. After that, the cells were resuspended in an unbuffered 100 mM NaCl solution and adjusted to an OD_600_ of 8.5. The measurements were performed in 3-ml aliquots of stirred cell suspension kept at ice-cold temperature (0.3–1.0 °C). The cells were illuminated using a halogen lamp, and the light-induced pH changes were monitored with a pH meter (Mettler Toledo). The measurements were repeated under the same conditions after the addition of 30 μM CCCP protonophore. For the measurements of the pH changes at low pH values, the pH of the final cell suspension was adjusted using 10% HCl solution.

### Cell culture and transfection

The patch-clamp recordings of CryoR1 were conducted in the neuroma glioblastoma cell line NG108-15 (ATCC, HB-12377TM, Manassas, USA) cultured in Dulbecco’s Modified Eagle Medium (DMEM, Sigma, St. Louis, USA) supplemented with 10 % fetal calf serum (Sigma, St. Louis, USA) and 1 % penicillin/streptomycin (Sigma, St. Louis, USA) (supplemented DMEM: DMEM+) at 37 °C and 5 % CO2. Cells were seeded on 24-well plates one day prior to transfection by Lipofectamine with pcDNA3.1(-) derivatives carrying the CryoR1 gene at a NG108-15 cell confluency of 50-70 %. For each well, a transfection mix of 100 µl DMEM, 2 µl Lipofectamine LTX (Invitrogen, Carlsbad, USA) and 500 ng of the plasmid DNA was prepared and added to a well with 400 µl of DMEM+. 24 h after transfection the medium was exchanged against 500 µl DMEM+ supplemented with 1 µM all-*trans* retinal.

### Manual patch-clamp recordings and data analysis

The electrophysiological characterization of CryoR1 was conducted by whole cell patch-clamp recordings of transiently transfected NG108-15 cells. Cells were patched two days after transfection under voltage-clamp conditions using the Axopatch 200B amplifier (Axon Instruments, Union City, USA) and the DigiData1440A interface (Axon Instruments, Union City, USA). Patch pipettes with a resistance of 2-6 MΩ were fabricated from thin-walled borosilicate glass on a horizontal puller (Model P-1000, Sutter Instruments, Novato, USA). The series resistance was <15 MΩ. The bath solution contained 140mM NaCl, 2mM CaCl_2_, 2mM MgCl_2_, 10mM HEPES, pH 7.4; and the pipette solution contained 110mM NaCl, 2mM MgCl_2_, 10mM EGTA, 10mM HEPES, pH 7.4. All recordings were performed at room temperature (20°C).

White light pulses were applied by a fast computer-controlled shutter (Uniblitz LS6ZM2, Vincent Associates, Rochester, USA) using Olympus U-HGLGPS 130W illumination system focused into an optic fiber (d=400 μm). The stationary photocurrents of the CryoR1 were measured in response to white light pulses with an intensity of 14 mW/mm^2^. The nanosecond light pulses of wavelengths ranging from 420 to 620 nm were applied using the the Opolette 355 tunable laser system (Opotek Inc, Carlsbad, USA) coupled into an optic fiber (d=400 μm). To activate the ground state, CryoR1 was preconditioned with 420 nm light pulses to ensure it was in the desired state. For the M2 state activation, prior stimulation was provided by 565 nm light pulses. This preparatory illumination, whether with 420 nm or 565 nm pulses, was continued until the protein no longer generated transient currents at a membrane potential of -40 mV.

The custom Python scripts developed in-house were used for data analysis. To calculate the absolute total charges transferred, we integrated the photocurrent over time for all sequential pulses until the photocurrents ceased. To determine the fractions of CryoR1 proteins in the two subpopulations, we compared the charge transfer in the first pulses recorded on the same cell at two different voltages. The ratio of charges transferred when activated with a 500 nm (620 nm) laser pulse at two different voltages is proportional to the fraction of the protein in the non-protonated (protonated) counterion subpopulation.

Illuminating with 420 nm light pulses returns all proteins to the ground state, implying that the total fractions (denoted as p) in the two subpopulations sum to 1 across all voltages:

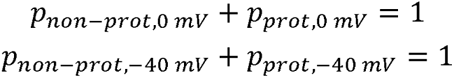

Assuming the total number of proteins remains constant during the experiment, the charge transferred (denoted as Q) after illumination with the first light pulse of a specified color is directly proportional to the fraction of proteins in the respective subpopulation:

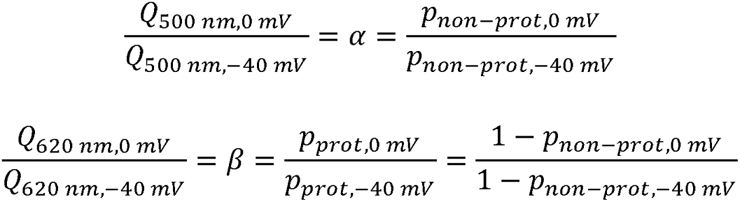

When combined, this allows us to derive the formula for the fraction of proteins in one of the subpopulations, which we used for our estimations:

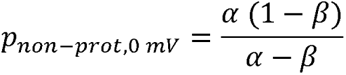

### Nanodisc reconstitution

For the dataset CryoR1 pH 8.0 in nanodisc, protein reconstitution was performed as previously described^64^. Namely, 20 mg/ml of liposomes containing *E.coli* polar lipids and egg PC (w/w 3:1, Avanti), were solubilized by adding 30 mM DDM followed by 1 min vortexing and 3 h incubation at 4 °C. Thawed purified rhodopsin, cleaved MSP2N2, and solubilized lipids were mixed in 3:5:100 molar ratio (considering a single protein chain) and incubated at 4 °C for 90 min while nutating. To remove the detergent, BioBeads (Bio-Rad) were added and the mixture was incubated overnight (10 hours) at 4 °C. After removing BioBeads with a syringe, the solution was supplied with 15 mM Imidazole, pH 8.0, and Ni-NTA resin, equilibrated with 50 mM Tris-HCl, pH 8.0, 100 mM NaCl to remove empty nanodiscs. After 1 h incubation at 4 °C on a rocking platform, resin was washed with 50 mM Tris-HCl, pH 8.0, 300 mM NaCl, 30 mM Imidazole, pH 8.0, and nanodiscs were eluted with 500 mM Imidazole in the same buffer. The sample was centrifuged (10 min, 20000g, 4 °C) and applied to Superdex 200 10/300 gel-filtration column (GE Healthcare) equilibrated with 20 mM Tris-HCl, pH 8.0, 100 mM NaCl. Fractions with nanodiscs were concentrated using VivaSpin® 500 MWCO 100 kDa concentrators (Sartorius) to the target concentration (around 2 mg/mL).

### Steady-state absorption spectroscopy, pH titration and illumination experiments

Absorption spectra of CryoRs samples were measured with an absorption spectrometer (Specord600, Analytik Jena). Before and after each experiment, absorption spectra were taken to check sample quality.

For the pH titration, equal amounts of the protein stock solution (3.3 µL) were added to 200 µL of the respective buffer to account for the same protein concentration in all prepared samples. The protein was suspended in the titration buffer containing 10 mM Trisodium citrate, 10 mM MES, 10 mM HEPES, 10 mM Tris, 10 mM CHES, 10 mM CAPS, and 10 mM Arg-HCl. The pH was adjusted with tiny amounts of [5000 mM] HCl or [5000 mM] NaOH, respectively.

For illumination experiments, solutions of each sample were prepared to have an optical density (OD) of ∼0.5 at an optical pathway of 10 mm. First of all, the dark-state spectrum was measured, second the main absorption band was illuminated using LEDs that match the absorption spectrum and in the final step the arised M-like state was illuminated. The following LEDs and LED settings were used: Thorlabs M420L3 (10 mA, 100 s), Thorlabs M530L (100 mA, 100 s), Thorlabs M565L2 (100 mA, 100 s), Thorlabs M590L3 (100 mA, 100 s), Thorlabs M625L2 (200 mA, 100 s).

Experiments at pH 3.5 were conducted in a 20 mM Sodium citrate buffer additionally consisting of 200 mM NaCl and 0.05% DDM. At pH 8.0, a buffer consisting of 20 mM Tris, 200 mM NaCl and 0.05% DDM was prepared. At pH 10.5, a 20 mM CAPS buffer with 200 mM NaCl and 0.05% DDM was used.

### Ultrafast transient absorption spectroscopy

Ultrafast transient absorption measurements were performed with a home-built pump-probe setup as described previously^65^. A Ti:Sa chirped pulse regenerative amplifier (MXR-CPA-iSeries, Clark-MXR Inc.) served as the fs-laser source and was operated at a central wavelength of 775 nm and with a repetition rate of 1 kHz, resulting in laser pulses with a pulse width of ∼150 fs. For excitation, the pulses from a two-stage NOPA setup were spectrally adjusted according to the absorption spectrum of the sample (600 nm and 620 nm for measurements at pH 3.5 and 560 nm for measurements at pH 10.5). For the probing of the photoinduced changes of sample absorbance, the laser fundamental was focused into a 5 mm CaF_2_-crystal to generate supercontinuum pulses. The generated white light was then split up and guided through sample and reference pathways, respectively. Two identical spectrographs (Multimode, AMKO), equipped with a grating (500 nm blaze, 1200 grooves per mm), a photodiode array (S8865-64, Hamamatsu Photonics) and a driver circuit (C9918, Hamamatsu Photonics) were used for signal detection.The obtained signals were digitized via a 16 bits data acquisition card (NI-PCI-6110, National Instruments). The pump and probe pulses were set to the magic angle (54.7°) configuration to eliminate anisotropic effects. Furthermore, the sample was constantly moved in a plane perpendicular to the excitation beam to avoid multiple excitation and sample degradation.

The samples were prepared to contain a protein concentration equal to an optical density of ∼0.3 for all measurements of CryoR1 at pH 3.5, ∼0.2 OD for the measurement of CryoR1 at pH 10.5 and ∼0.15 OD for the measurements of CryoR2 at both pH 3.5 and pH 10.5. Due to their slow photocycle kinetics at pH 10.5, respective samples were constantly illuminated using a Thorlabs M405L4 LED, operated at 300 mA.

### Transient flash photolysis spectroscopy

A Nd:YAG laser (SpitLight 600, Innolas Laser) was used to pump an optical parametric oscillator (preciScan, GWU-Lasertechnik). The OPO was set to generate excitation pulses according to the absorption maximum of the respective sample at an average pulse energy of ∼2.2 mJ/cm^2^. A Xenon or a Mercury-Xenon lamp (LC-8, Hamamatsu) served as probe light sources. Two identical monochromators (1200 L/mm, 500 nm blaze), one in front and one after the sample, set the chosen probing wavelengths. Absorption changes were detected by a photomultiplier tube (Photosensor H6780-02, Hamamatsu) and then converted into an electrical signal, which was recorded by two oscilloscopes (PicoScope 5244B/D, Pico Technology) with overlapping timescales. For each transient 30 acquisitions were measured and averaged to increase the S/N ratio. To obtain data files with a reasonable size for further analysis, raw data files were reduced using forward averaging and a combined linear and logarithmic timescale.

Detergent-solubilized samples were measured in a 2 x 10 mm quartz cuvette and prepared to have a protein concentration equal to an optical density of ∼1.0 at an optical path length of 10 mm. For the measurement presented in Fig. 3B, a Thorlabs M405L4 LED operated at 300 mA repopulated the parent state.

For the measurement of CryoR1 at pH 3.5, the time point at 468.2 ms was excluded from the dataset after data reduction due to an artifact from the coupling of the two oscilloscopes.

### Analysis of time-resolved spectroscopic data

Analysis of time-resolved spectroscopic data was performed using OPTIMUS software^66^. The data of the ultrafast transient absorption and transient flash photolysis measurements were objected to the model-free lifetime distribution analysis (LDA) yielding the lifetime distributions of the individual photointermediate transitions, which are summarized in a lifetime density map (LDM). The mentioned lifetimes of the photointermediate transitions were obtained from the point with maximum amplitude of the respective lifetime distribution. For the flash photolysis measurement of CryoR1 at pH 10.5, a global lifetime analysis (GLA) was performed, yielding decay-associated spectra (DAS) and the lifetimes of the respective photointermediate transitions.

### Cryo-EM grid preparation and data collection

All samples were concentrated to 30-50 mg/ml using 100,000 MWCO concentrators (Millipore) at pH 8.0 and later mixed with a buffer with pH 8.0 (both pH 8.0 structures) or pH 4.3 (pH 4.3 structure). For grids at pH 4.3, sample was additionally concentrated to 30 mg/ml after adding buffer, and mixed again with pH 4.3 buffer, to remove excess of pH 8.0 buffer. After that, all samples were diluted (to 7 mg/ml for CryoR2 dataset, and CryoR1 datasets at pH 10.5 M and ground state; to 10 mg/ml for CryoR1 at pH 4.3 and 8.0, and to 2.7 mg/ml for CryoR1 in nanodiscs), and volume of applied onto freshly glow-discharged (30s at 5 mA) Quantifoil grids (Au R1.2/1.3, 300 mesh) at 20 °C and 100% humidity and plunged-frozen in liquid ethane. The cryo-EM data were collected using either 300[keV Krios microscope (Thermo Fisher), equipped with Gatan K3 detector (all dataset except for CryoR1 at pH 8.0), or 200 keV Talos Arctica (Thermo Fisher), equipped with Gatan K2 Summit detector (CryoR1 at pH 8.0).

### Cryo-EM data processing

All steps of data processing were performed using cryoSPARC v.4.0.2^67^ (Fig. S17). Motion correction and contrast transfer function (CTF) estimation were performed with default settings for all datasets. For datasets CryoR1 pH 8.0, pH 10.5 ground, and pH 10.5 M state, particle picking was performed using Topaz^68^ pre-trained model, followed by duplicate removal with 50 Å distance. For datasets CryoR1 pH 4.3 and pH 8.0, initial volumes were generated after picking with Topaz and blob picking, respectively, followed by template picking using generated volumes.

For datasets CryoR1 pH 10.5 ground and M states, pH 8.0 in nanodiscs, and CryoR2 pH 8.0, picked particles were extracted with 3-5x binning (up to 128 px). An initial set of particles was cleaned using two rounds of 2D classification (first round: 80 classes, 80 iterations, batch size 200-400, use clamp-solvent: true; second round: 20-40 classes, 40 iterations, batch size 50-200, use clamp-solvent: true). After that, particles were cleaned using a "3D classification" (ab initio model generation with 5 classes, followed by heterogeneous refinement; in case of CryoR1 pH 8.0 in nanodisc, 2 cycles of “3D classification” were performed, see Fig. S17). These particles were re-extracted without binning, followed by non-uniform (CryoR2 and CryoR1 pH 10.0 ground state) or homogeneous (CryoR1, pH 8.0 in nanodisc, pH 10.5 M state) refinement (C5 symmetry, with per-particle CTF and defocus refinement) and local (all but pH 10.5 M state) refinement (C5 symmetry), yielding final maps.

For datasets CryoR1 pH 4.3 and pH 8.0, particles were extracted using 2x binning (pH 4.3) or no binning (pH 8.0). An initial set of particles was cleaned using one round of 2D classification with default parameters (pH 4.3) or two rounds of 2D classification (first iteration: use clamp-solvent=true, second iteration: batchsize 200, 20 classes). These particles were subjected to *ab initio* model generation with 2 classes, for pH 4.3 followed also by an unbinned particle re-extraction. After that, particles were refined with non-uniform refinement (C5 symmetry, with per-particle CTF and defocus refinement) and local refinement (C5 symmetry, mask=dynamic for pH 8.0), yielding final maps.

### Model building and refinement

The pentameric model of CryoR1 was generated using Alphafold^69^ and docked as a rigid body into cryo-EM maps manually in ChimeraX. Further refinement was performed using Phenix^70,71^ and Coot^72^, producing the final statistics described in Table S3. Visualization and structure interpretation were carried out in UCSF Chimera^73,74^ and PyMol (Schrödinger, LLC).

### Crystallization

The crystals of CryoR2 were grown with an *in meso* approach^75^, similar to that used in our previous works^45,46^. In particular, the solubilized protein (80 mg/ml) in the crystallization buffer was mixed with octyl-β-D-Glucopyranosid (OG, Glycon) and then with the premelted at 42°C monoolein (MO, Nu-Chek Prep) in a 2:1 ratio (protein/OG:lipid) to form a lipidic mesophase. The mesophase was homogenized in coupled syringes (Hamilton) by transferring the mesophase from one syringe to another until a homogeneous and gel-like material was formed.

Then, 150 nl drops of a protein–mesophase mixture were spotted on a 96-well LCP glass sandwich plate (Marienfeld) and overlaid with 400 nL of precipitant solution by means of the Mosquito crystallization robot (SPT Labtech). Crystals were obtained with the final protein concentration of 40 mg/ml and the final OG concentration of 9% (w/v) in the water part of the mesophase. The best crystals were obtained using 1.2M Na/K-Pi pH 4.6 as a precipitant. The crystals were grown at 22°C and appeared in 1 month.

For the determination of the CryoR2 crystal structure, once the crystals reached their final size, crystallization wells were opened, and drops containing the protein-mesophase mixture were covered with 100 μl of the precipitant solution. For the data collection, harvested crystals were incubated for 5 min in the precipitant solution. Crystals were harvested using micromounts (Mitegen, USA), flash-cooled, and stored in liquid nitrogen.

### Time-resolved microspectrophotometry on the CryoR2 crystals

The spectroscopic characterization of the CryoR2 crystals was performed at the *ic*OS Lab of the ESRF^76^ using the TR-*ic*OS instrument^77^. Measurements were conducted at room temperature with crystals of 100 x 20 x 15 µm^3^ mounted in a dedicated holder. Series of transient UV-Vis absorption spectra were measured based on a pump-probe scheme. A nanosecond laser (Surelite EX, Amplitude Technologies, France), combined with an optical parametric oscillator (OPO) (Horizon II, Amplitude Technologies, France), was used to produce the pump signal. Laser pulses were collimated in a dedicated laser optical box into a 910 µm diameter high-energy optical fiber (Thorlabs, USA) connected to a parabolic mirror on the top of the optomechanical setup of the TR-*ic*OS instrument. A 15x reflective objective is used to focus light at the sample position, with a ∼130 µm focal spot (1/e^2^ cutoff). The probe signal was provided by a xenon flash lamp module (L11316-11, Hamamatsu Photonics, Japan) connected via a 400 µm diameter optical fiber (Avantes, The Netherlands) to a second parabolic mirror positioned to match the optical path of the pump signal. Focusing of the probe light is achieved through the same 15x reflective objective, resulting in a ∼65 µm focal spot (1/e^2^ cutoff) at the sample position. Two different wavelengths were used for the pump signal, 532 and 630 nm, with fluences of 170 mJ/cm^2^ and 151 mJ/cm^2^ respectively. Delays between pump and probe signals were controlled using the ESRF CITY timing module and varied between 10 µs and 10 s. Spectra were measured using a CMOS spectrophotometer (AvaSpec-ULS2048CL-EVO-RS-UA, Avantes, The Netherlands), with an integration time of 100 µs. In-house Python script was used to plot the data (https://github.com/ncara/TRicOS).

### Accumulation of the intermediate state in CryoR2 crystals

For the accumulation and cryotrapping of the M_2_ state, the crystal was originally kept at 100 K. It was then illuminated by a 532-nm-laser with the nitrogen stream simultaneously blocked for 2 s. Due to the short illumination time the crystal was not rotated during the illumination. For better illumination, the crystal was oriented with the largest plane perpendicular to the laser beam. The laser was then switched off once the crystal was back at 100 K. The laser power density of 10 W/cm^2^ at the position of the sample was used. The laser power was adjusted to maximize the fraction of the accumulated M_2_ state; the absence of destructive effect of the laser on crystals was justified by the remaining diffraction quality after the cryotrapping procedure.

The mean size of the crystals was 100×20×20 μm^3^. The rod-shaped crystals were oriented so that the one of the largest planes (100×20 μm^2^) was perpendicular to the laser beam. The laser beam was focused to the size of 200×200 μm^2^ (1/e^2^).

### Diffraction data collection and treatment

X-ray diffraction data were collected at the P14 beamline of PETRAIII (Hamburg, Germany) using an EIGER2 X 16M CdTe detector. The data collection was performed using MxCube2 software. Diffraction images were processed using XDS^78^. The reflection intensities were scaled and merged using the Staraniso server^79^. There is no possibility of twinning for the crystals. Diffraction data from a single crystal were used. The data collection and treatment statistics are presented in Table S4.

### Crystal structure determination and refinement

Initial phases were successfully obtained in the C2221 and C121 space groups by molecular replacement using MOLREP^80^ from the CCP4 program suite^81^ using the cryo-EM structure of CryoR2 as a search model. The initial model was iteratively refined using REFMAC5^82^ and Coot^83^. The structure refinement statistics are presented in Table S2. The difference F_olight_-F_odark_ were built using the PHENIX program suite.

## DATA AVAILABILITY

Atomic models built using X-ray crystallography and cryo-EM data have been deposited in the RCSB Protein Data Bank with PDB codes 8R96 (crystal structure of CryoR2 in the dark state, type A crystals), 8R97 (crystal structure of CryoR2 in the dark state, type B crystals), 8R98 (crystal structure of CryoR2 in the illuminated state, type B crystals), 8R0K (cryo-EM structure of CryoR1 at pH 4.3 in detergent), 8R0L (cryo-EM structure of CryoR1 at pH 8.0 in nanodisc), 8R0M (cryo-EM structure of CryoR1 at pH 8.0 in detergent), 8R0N (cryo-EM structure of CryoR1 at pH 10.5 in detergent, the ground state), 8R0O (cryo-EM structure of CryoR1 at pH 10.5 in detergent, the M state), and 8R0P (cryo-EM structure of CryoR2 at pH 8.0 in detergent). The cryo-EM density maps have been deposited in the Electron Microscopy Data Bank under accession numbers EMD-18795 (cryo-EM structure of CryoR1 at pH 4.3 in detergent), EMD-18796 (cryo-EM structure of CryoR1 at pH 8.0 in nanodisc), EMD-18797 (cryo-EM structure of CryoR1 at pH 8.0 in detergent), EMD-18798 (cryo-EM structure of CryoR1 at pH 10.5 in detergent, the ground state), EMD-18799 (cryo-EM structure of CryoR1 at pH 10.5 in detergent, the M state), and EMD-18800 (cryo-EM structure of CryoR2 at pH 8.0 in detergent).

## ACKNOWLEDGEMENTS

We are deeply thankful to Lorna Richardson and Martin Beracochea for the help with the analysis of the genes retrieved from the MGnify database. We are also deeply thankful to Dr. Andrey Rozenberg for his comments on the evolutionary affiliation of CryoRhodopsins. We are deeply thankful to Prof. Dr. Ernst Bamberg for the discussion of the electrophysiology and potential optogenetic applications of CryoRhodopsins. This research was supported by the German Research Foundation (CRC 1507 – Membrane-associated Protein Assemblies, Machineries and Supercomplexes; Project 05 to J.W.). This work used the *ic*OS platform of the Grenoble Instruct-ERIC center (ISBG; UAR 3518 CNRS-CEA-UGA-EMBL) within the Grenoble Partnership for Structural Biology (PSB), supported by FRISBI (ANR-10-INBS-0005-02) and GRAL, financed within the University Grenoble Alpes graduate school (Ecoles Universitaires de Recherche) CBH-EUR-GS (ANR-17-EURE-0003). E.M., A.S. and A.G. thank NeCEN personnel for the support during data collection. The access to NeCEN facilities was funded by the Netherlands Electron Microscopy Infrastructure (NEMI), project number 184.034.014 of the National Roadmap for Large-Scale Research Infra-structure of the Dutch Research Council (NWO). A.G. was supported by NWO grant OCENW.KLEIN.141. K.K. has been supported by EMBL Interdisciplinary Postdoctoral Fellowship (EIPOD4) under Marie Sklodowska-Curie Actions Cofund (grant agreement number 847543). The work of A.A. was supported by funding from the German Research Foundation to T. Moser via the Multiscale Bioimaging - Cluster of Excellence (EXC 2067/1-390729940).

## AUTHOR CONTRIBUTIONS

K.K. performed bioinformatics analysis and identified CryoRhodopsins group under the supervision of A.B.; K.K. did cloning and protein production; K.K. did functional tests in *E.coli* cell suspension; A.A. did cloning of the CryoR1 for the mammalian expression; A.A. and T.Mager. planned the patch-clamp electrophysiology experiments; A.A. performed electrophysiology experiments and analyzed the data; A.A. and T.Mager. wrote the text for the electrophysiology part; G.H.U.L., A.V.S., and M.Asido. did the spectroscopy experiments under the supervision of J.W.; K.K. crystallized CryoR2, collected and processed X-ray crystallography data, and solved the crystal structure of CryoR2; G.B. helped with the X-ray crystallography data analysis; S.R., S.E., and K.K. performed time-resolved microspectrophotometry on the CryoR2 crystals; N.C. did the time-resolved microspectrophotometry data processing and visualization; A.R. supervised the time-resolved microspectrophotometry experiments; A.S. prepared grids and performed initial characterization and analysis of EM data; K.K., E.M., and A.S. did the preillumination of the samples for cryo-EM grid preparation and controlled the illumination conditions during the grid preparation; M.Agthe. prepared the LED and laser light sources for the illumination of CryoRhodopsins and installed the optical system for the intermediate state cryotrapping at the P14 station; E.M. assembled nanodiscs with CryoR1; E.M. and A.G. processed cryo-EM data and obtained initial pentameric models of CryoR1 and CryoR2; K.K. refined the cryo-EM structures; A.G. supervised all steps of cryo-EM pipeline and structure refinement and validation; K.K., G.H.U.L., E.M. analyzed the data and prepared a manuscript with contributions of V.B., A.G., G.B., T.R.S., A.B., J.W., A.A., T.Mager., T.Moser. and all other authors.

## COMPETING INTEREST

Authors declare no competing interests.

## SUPPLEMENTARY INFORMATION

### Supplementary Figures

**Fig. S1.**
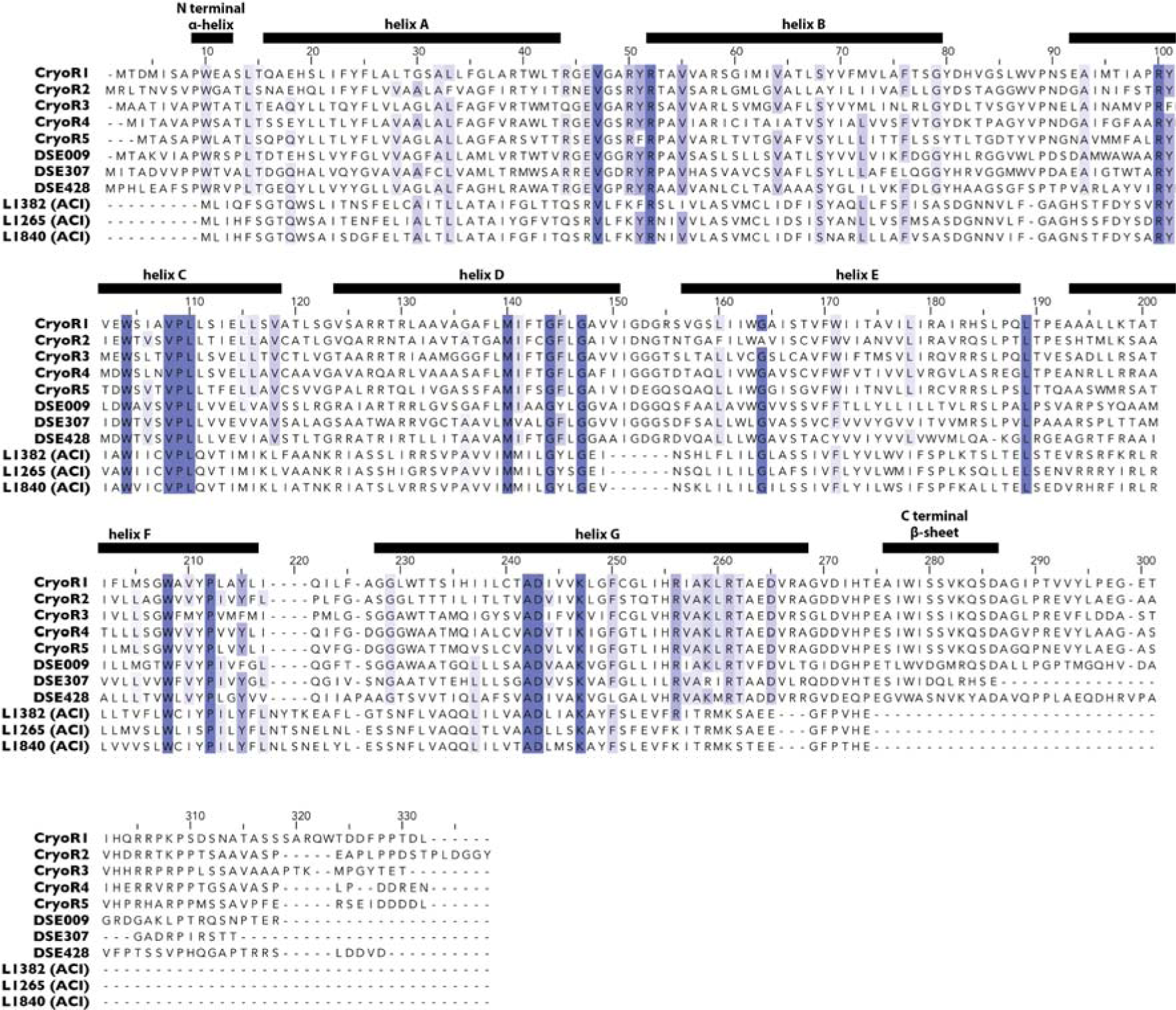
Multiple sequence alignment of CryoRs, DSE, and ACI rhodopsins.

**Fig. S2.**
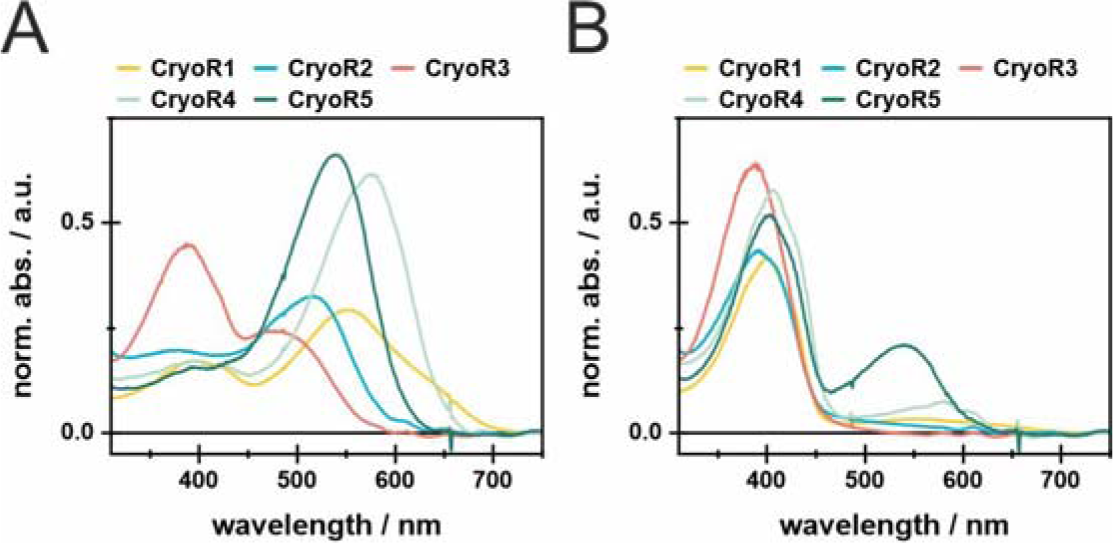
Dark state and PSS spectra for CryoR1-5 at pH 8.0. **A.** dark state spectra of all investigated CryoRs at pH 8.0. **B.** spectra of the populated PSS for all investigated CryoRs after illumination of the main absorption band in the range of 530 nm to 590 nm.

**Fig. S3.**
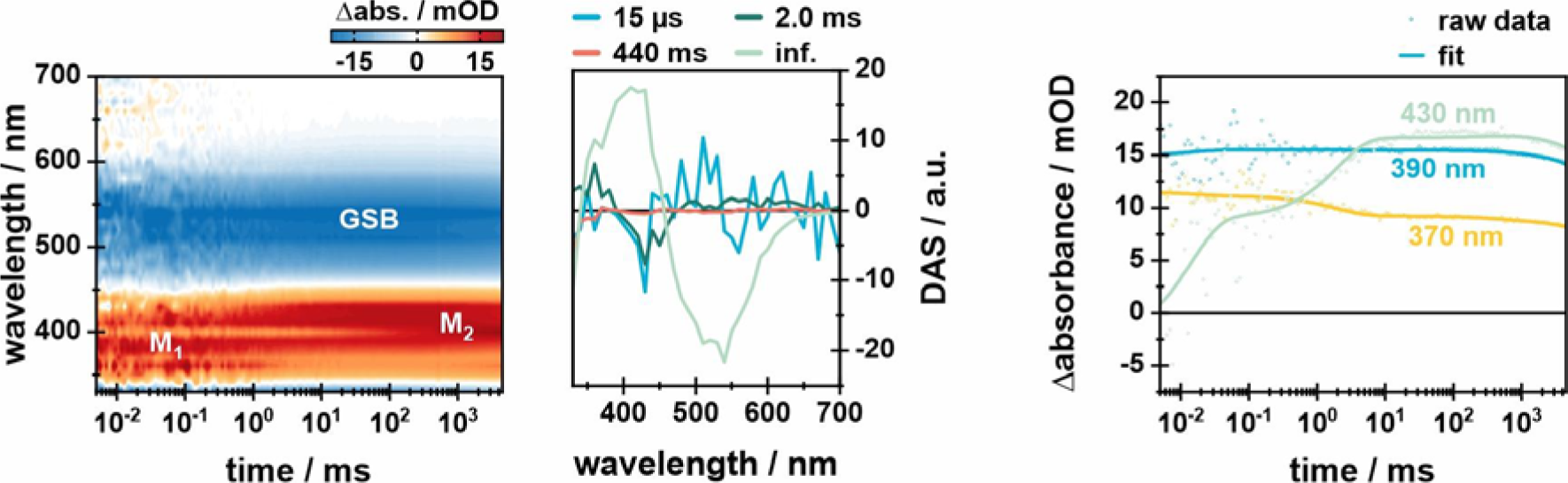
Flash-photolysis data of CryoR1 showing the intermediates M_1_ and M_2_. Flash-photolysis measurement of CryoR1 at pH 10.5 is shown as a 2D-contour plot. The photocycle was measured until 4.5 s and ended afterwards via illumination with a 405 nm LED (300 mA) before each acquisition. Additionally, the decay-associated spectra (DAS) are shown retrieving the kinetic information of the measurement as well as the lifetimes of the observed photointermediate transitions. Furthermore, transients at three characteristic wavelengths (370 nm, 390 nm, and 430 nm) are shown in the right panel (raw data as dots, fit as lines).

**Fig. S4.**
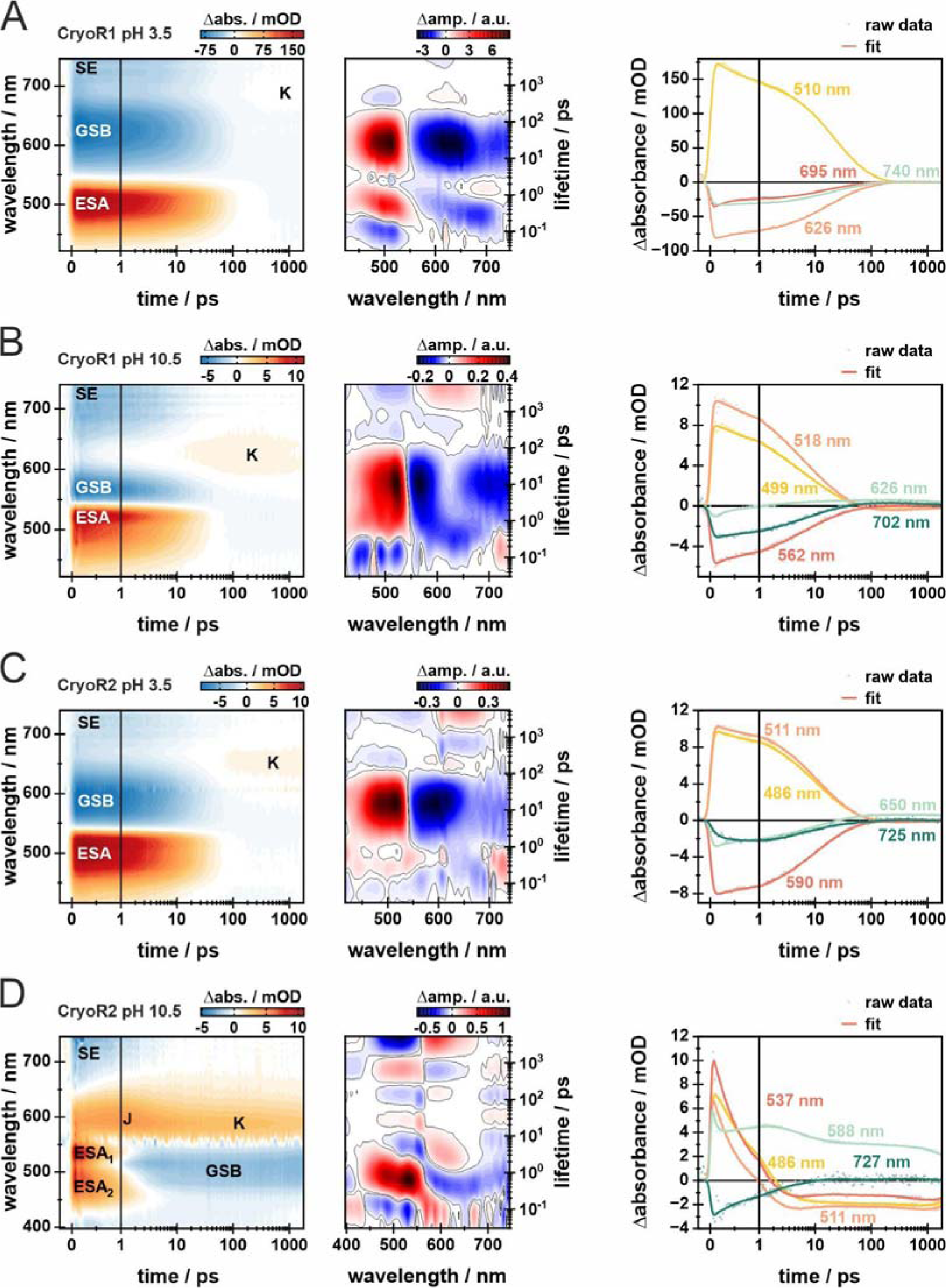
Ultrafast dynamics of CryoR1 and CryoR2 at low and high pH. Ultrafast dynamics of CryoR1 at **A** pH 3.5 and **B** pH 10.5 are displayed together with the ultrafast dynamics of CryoR2 at **C** pH 3.5 and **D** pH 10.5 as 2D-contour plots. The x-axis is linear until 1 ps and logarithmic afterward. The signal amplitude is color coded as follows: positive (red), no (white) and negative (blue) Δabs. Additionally, the corresponding lifetime density maps (LDM) are shown illustrating the kinetics components of the respective measurement. Furthermore, transients at specific wavelengths are shown. The raw data is shown as dots, while the obtained fit is shown as lines.

**Fig. S5.**
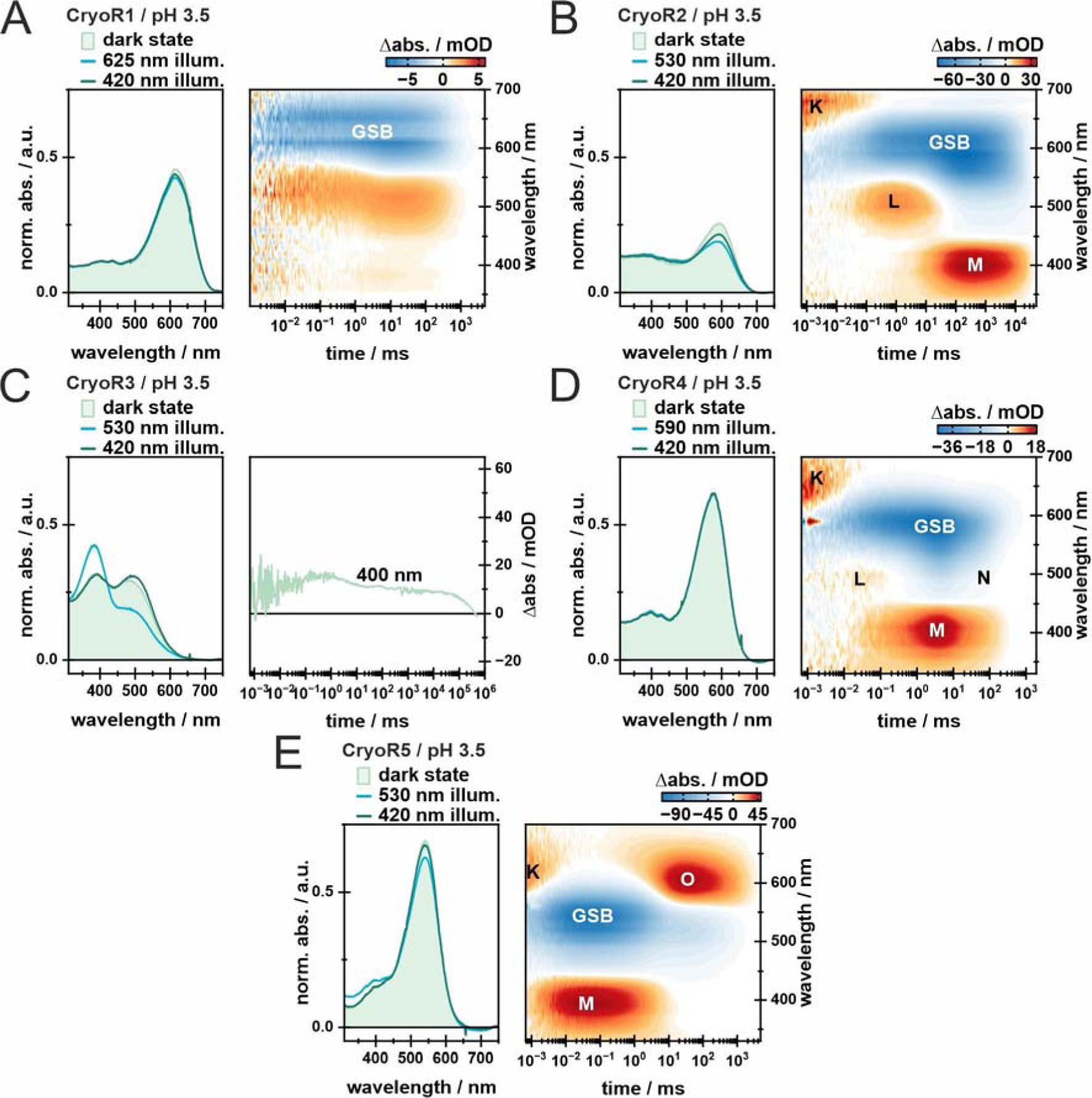
Illumination experiments and photocycle kinetics of CryoRs at low pH. In each panel, the dark state spectra (light green), as well as the PSS spectra after illumination of the main absorption band (blue) and the spectra after illumination of the obtained PSS with blue light (dark green) to recover the dark state of each investigated cryoR are shown. Each LED was turned on for 100 s. LEDs in the 500 nm range were operated at 100 mA, while the 420 nm LED was operated at 10 mA. The 625 nm LED was operated at 200 mA. Additionally, for all investigated CryoRs except CryoR3 the measured photocycle kinetics were depicted as 2D-contour plots. The signal amplitude is color coded as follows: positive (red), no (white) and negative (blue) Δabs. The observed photocycle intermediates are named in the common way for microbial rhodopsins based on the photocycle of bacteriorhodopsin. Due to the slow photocycle kinetics, only the 400 nm transient was measured and is shown for CryoR3, similar to the measurements at high pH shown in Fig. 2 in the main text.

**Fig. S6.**
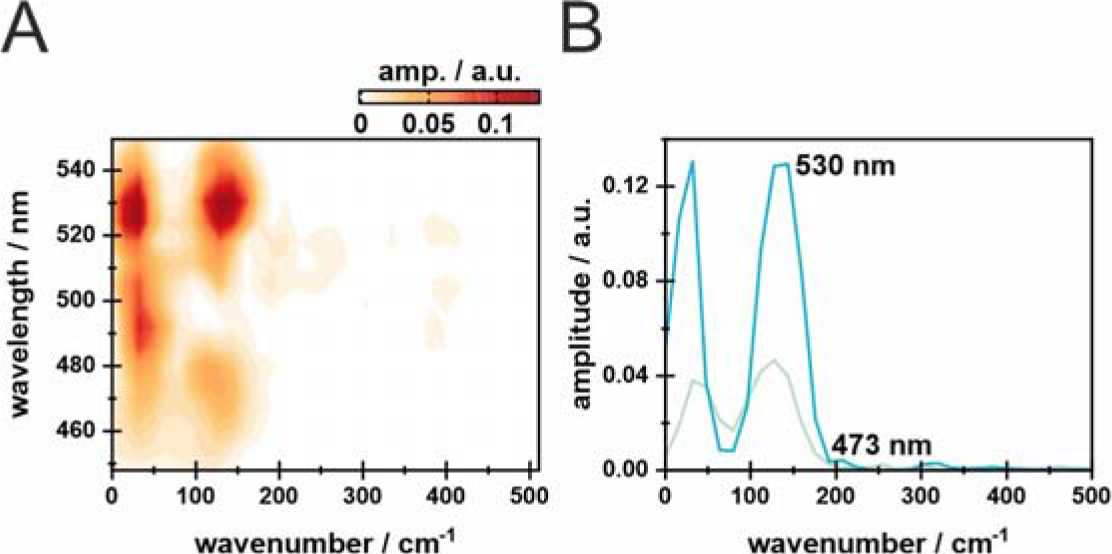
Fourier analysis of the observed coherent oscillations in CryoR1 at pH 3.5. **A** shows the Fourier-transformed spectra of CryoR1 at pH 3.5 as a 2D-contour plot. Areas with a high amplitude show frequency contributions of the observed coherent oscillations. **B** shows the Fourier-transformed spectra at two specific wavelengths to further elucidate the frequencies of the observed oscillations.

**Fig. S7.**
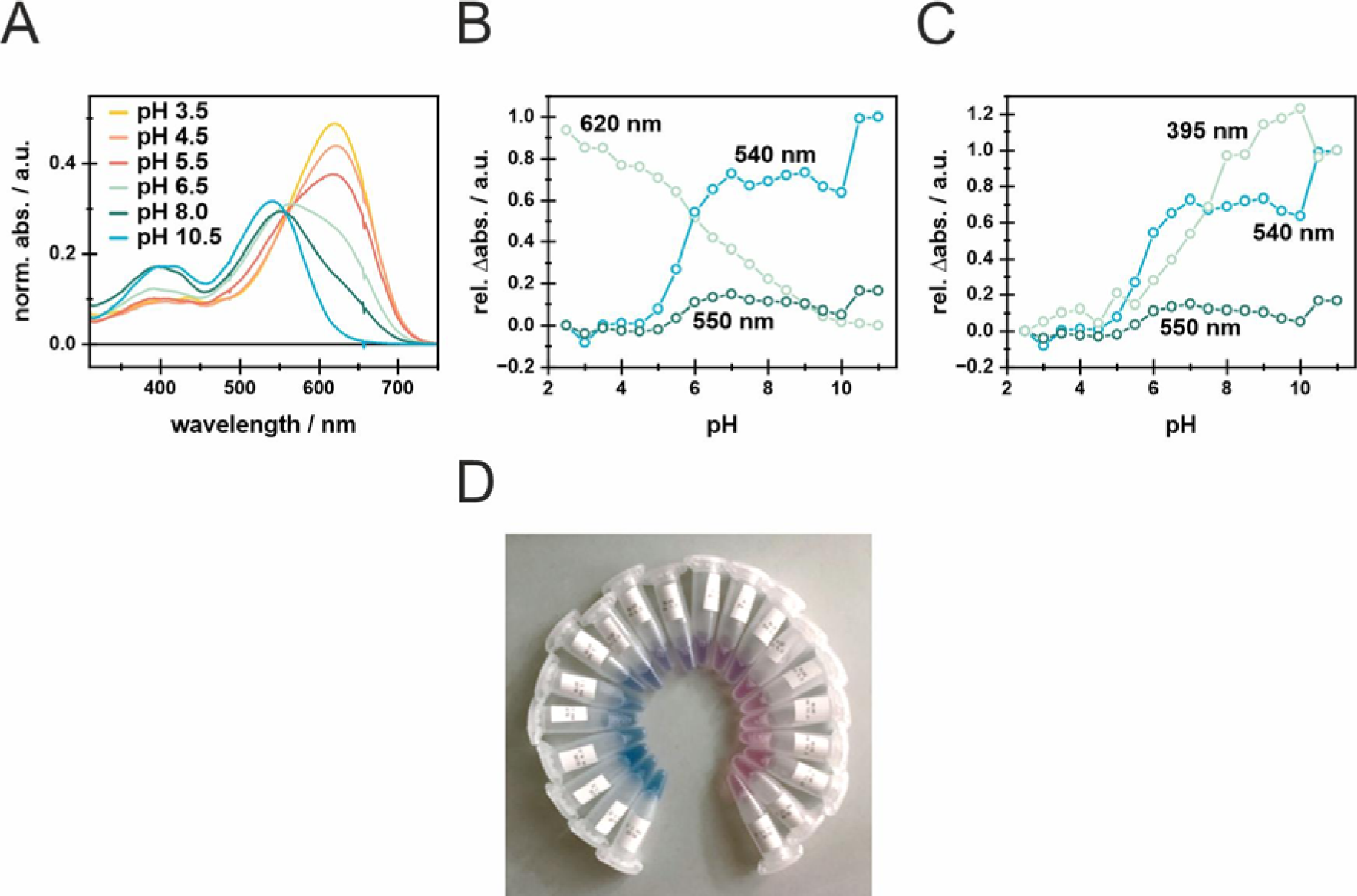
pH titration of CryoR1 in the range of pH 2.5 to pH 11.0. **A** absorption spectra at specific pH values indicating the observed pH-dependent shift of the absorption maximum. Furthermore, the change in absorption is shown in a pH-dependent manner for **B** 540 nm, 550 nm and 620 nm and **C** 395 nm, 540 nm and 550 nm. The absorption changes are shown as relative absorption changes. For 620 nm it was assumed that the maximum amplitude is reached at pH 2.5 and therefore the obtained value was set to 1. For all other shown wavelengths, it was assumed that the maximum amplitude is reached at pH 11.0, so the rel. Labs. was set to 0 at pH 2.5. **D** shows the samples used to measure the absorption spectra for the pH titration in the range of pH 2.5 (left) to pH 12.0 (right).

**Fig. S8.**
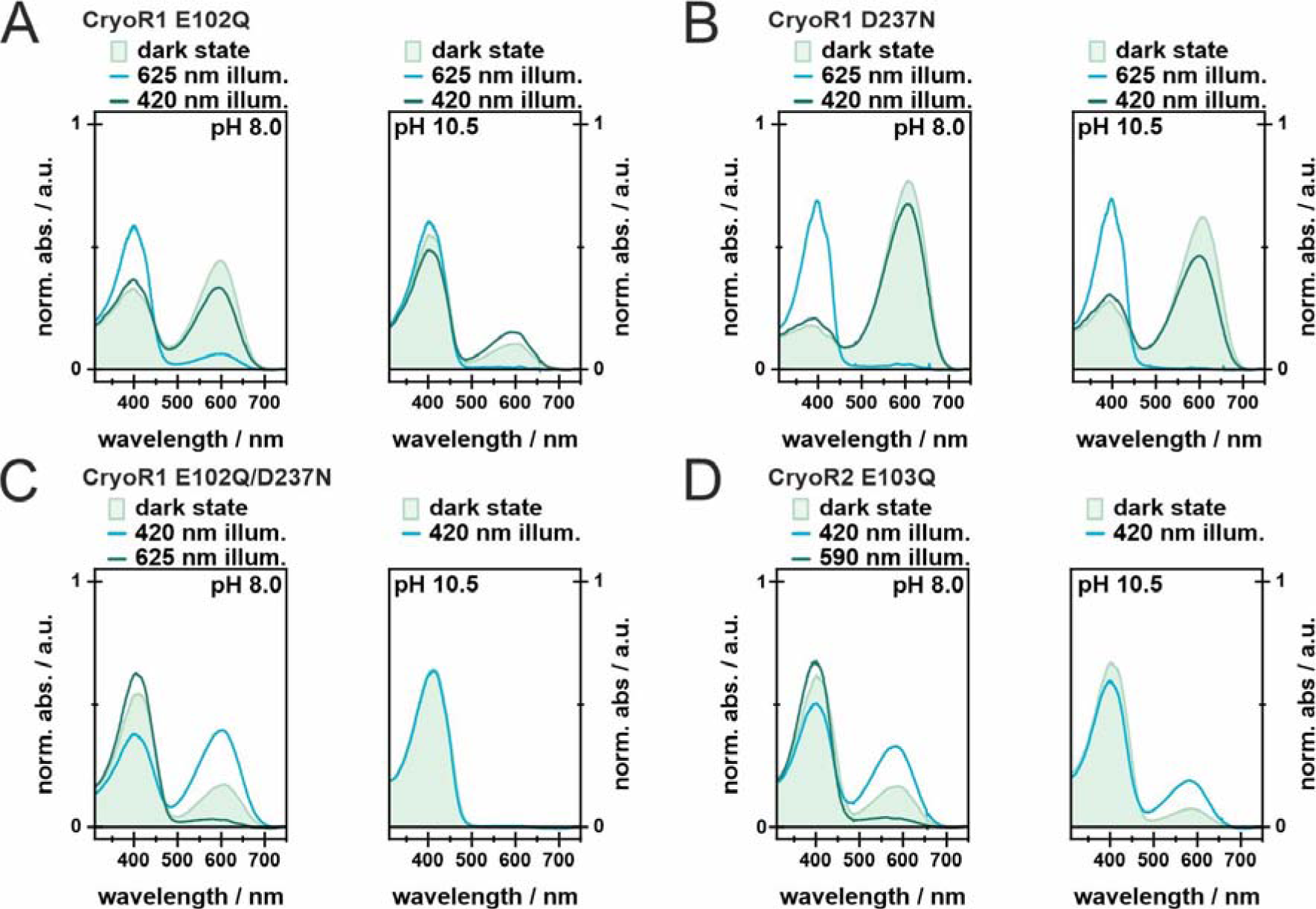
Illumination experiments of mutants of charged residues of CryoR1 and CryoR2 in their retinal binding pocket. Dark state and PSS spectra of CryoR1 mutants E102Q **A**, D237N **B,** and E102Q/D237N **C**, as well as CryoR2 E103Q **D**. First of all, the dark state spectrum was measured, followed by the PSS after illumination of the main absorption band for 100s and the illumination of the potentially rising band for 100s afterward. The 625 nm LED was operated at 200 mA, the 590 nm LED was operated at 100 mA and the 420 nm LED was operated at 10 mA.

**Fig. S9.**
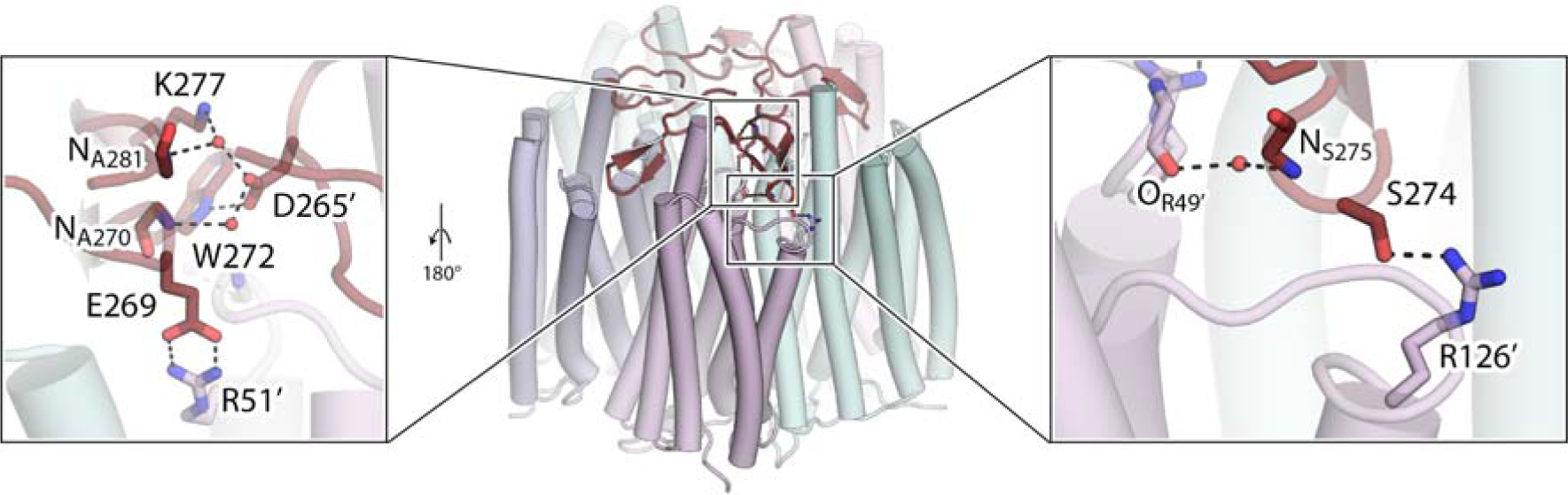
Involvement of the C terminus in stabilizing the CryoR1 pentamer. The overall view of the pentamer of CryoR1 (center) with the C terminus colored red. Detailed view of the interprotomeric contacts formed by the tip of the β-sheet of the C terminus (colored red) with the nearby protomer (colored light purple) (right). Detailed view of the interprotomeric contacts formed by the C terminus in the region closer to the central axis of the pentamer (left). For clarity, the view in the left panel is from the center of the pentamer.

**Fig. S10.**
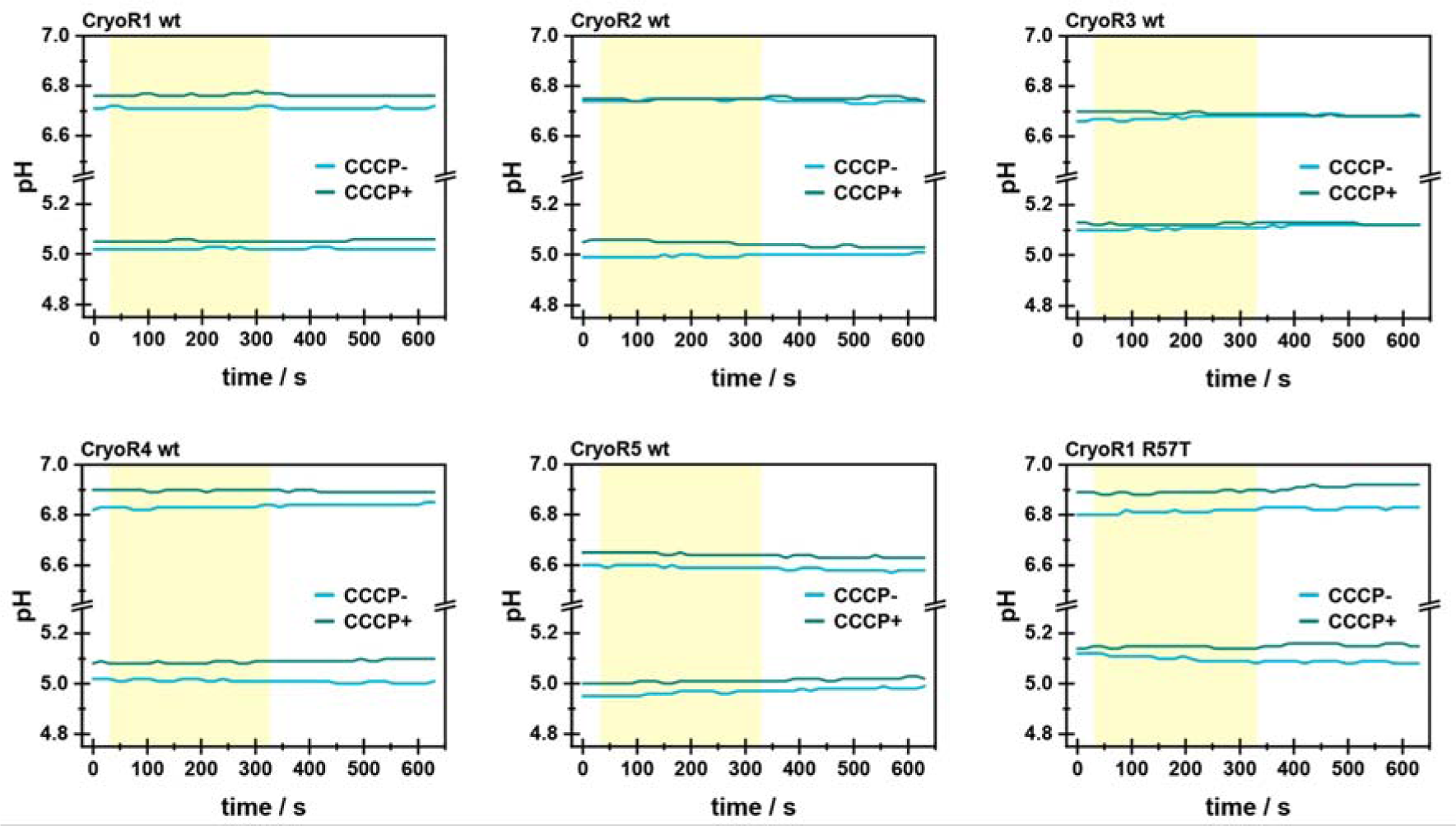
Functional tests of CryoRs in *E.coli* cell suspension. Light-induced pH changes in a suspension of *E. coli* expressing CryoRs in non-buffered 100 mM NaCl solution. Experiments were performed at two pH values; pH was adjusted using 10% HCl. Cyan lines represent results from experiments without any protonophore; green lines show pH changes in the presence of 30 μM of protonophore carbonyl-cyanide m-chlorophenyl-hydrazone (CCCP). The yellow area indicates the period of time in which the white light was on.

**Fig. S11.**
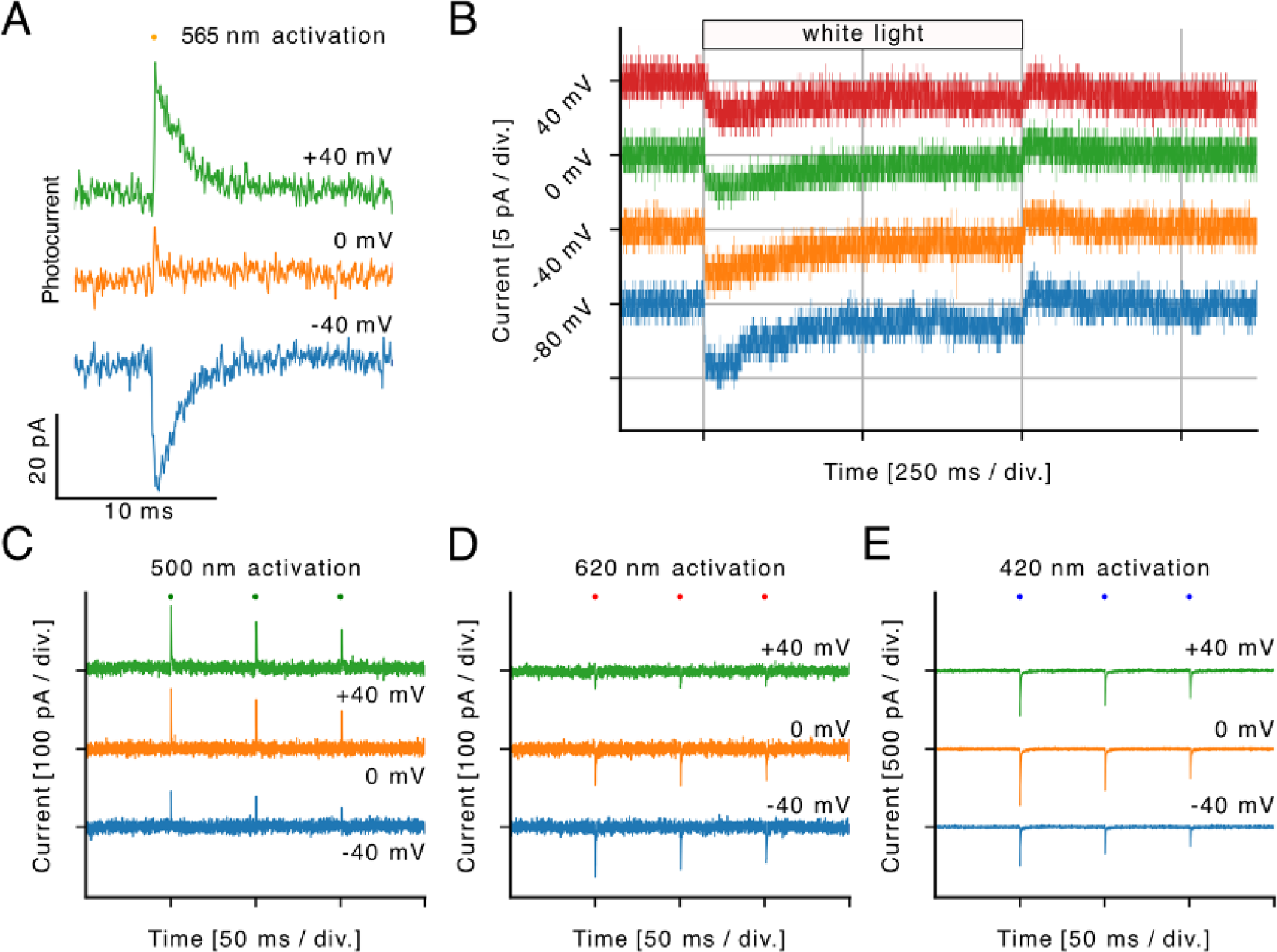
Electrophysiological characterization of CryoR1 in NG108-15 cells. **A.** Voltage-dependence of CryoR1 photocurrents elicited by nanosecond laser pulses with a wavelength of 565 nm. **B.** Stationary CryoR1 photocurrents upon illumination with white light. **C-E.** CryoR1 photocurrents at the shown voltages elicited by nanosecond laser pulses with a wavelength of **(C)** 500 nm for the activation of the ground state subpopulation with the deprotonated counterion complex, **(D)** 620 nm for the activation of the ground state subpopulation with the protonated counterion complex, and **(E)** 420 nm for M_2_ state activation.

**Fig. S12.**
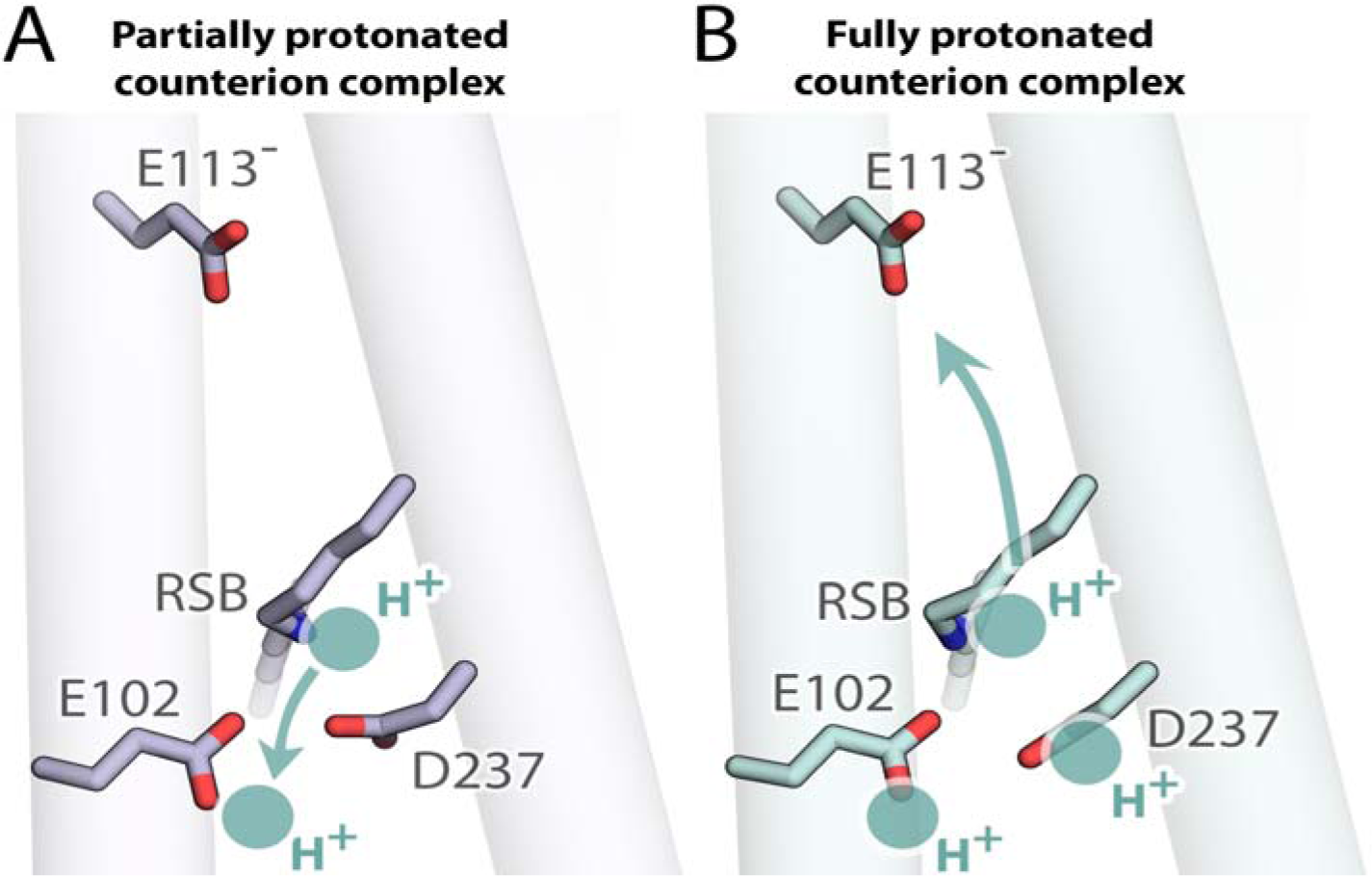
Proposed scheme of the proton transfer in CryoR1. **A.** The RSB proton transfer towards the partially protonated counterion complex upon activation with the 500 nm laser pulses. B. The RSB proton transfer towards the cytoplasmic side to the deprotonated E113 residue upon illumination with the 620 nm laser pulses when the counterion complex is fully protonated.

**Fig. S13.**
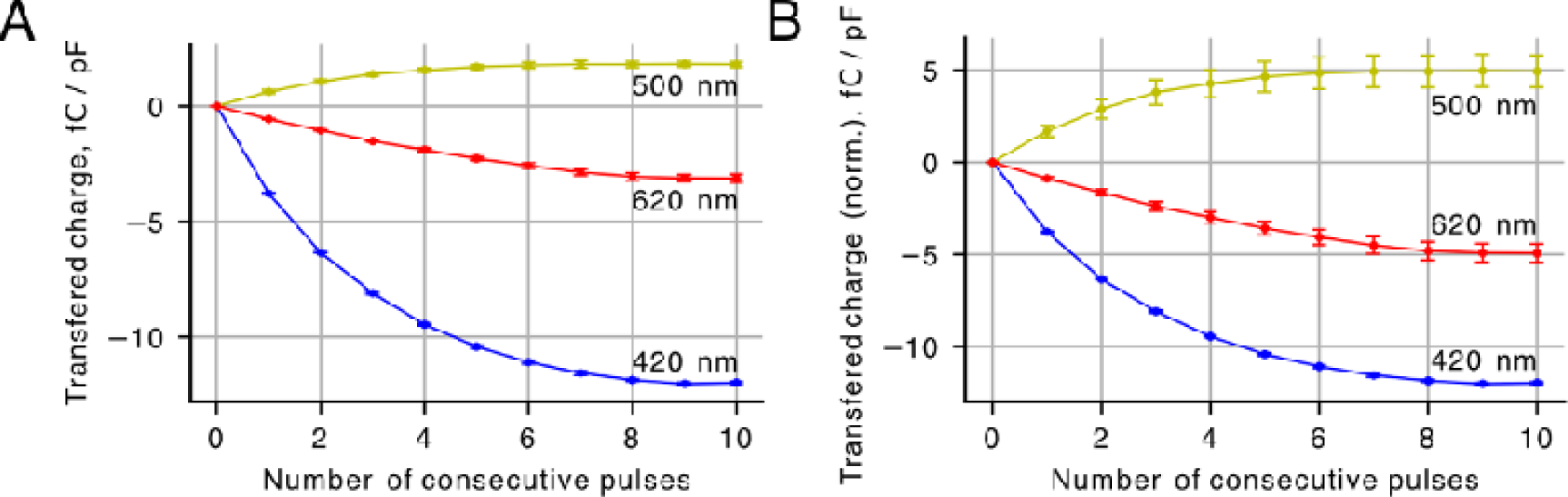
Quantification of the transferred charge. **A.** Cumulative charge transferred at a membrane voltage of 0 mV in response to illumination with consecutive light pulses of 420 nm (for the M_2_ state activation), 500 nm, and 620 nm, normalized by cell capacitance (number of cells, N = 2). **B.** Cumulative charge transferred at a membrane voltage of 0 mV in response to illumination with consecutive light pulses of 420 nm (for the M_2_ state activation), 500 nm, and 620 nm, normalized by cell capacitance. The curves for 500 nm and 620 nm activation were further normalized by the fractions of the protein in the respective subpopulations (number of cells, N = 2).

**Fig. S14.**
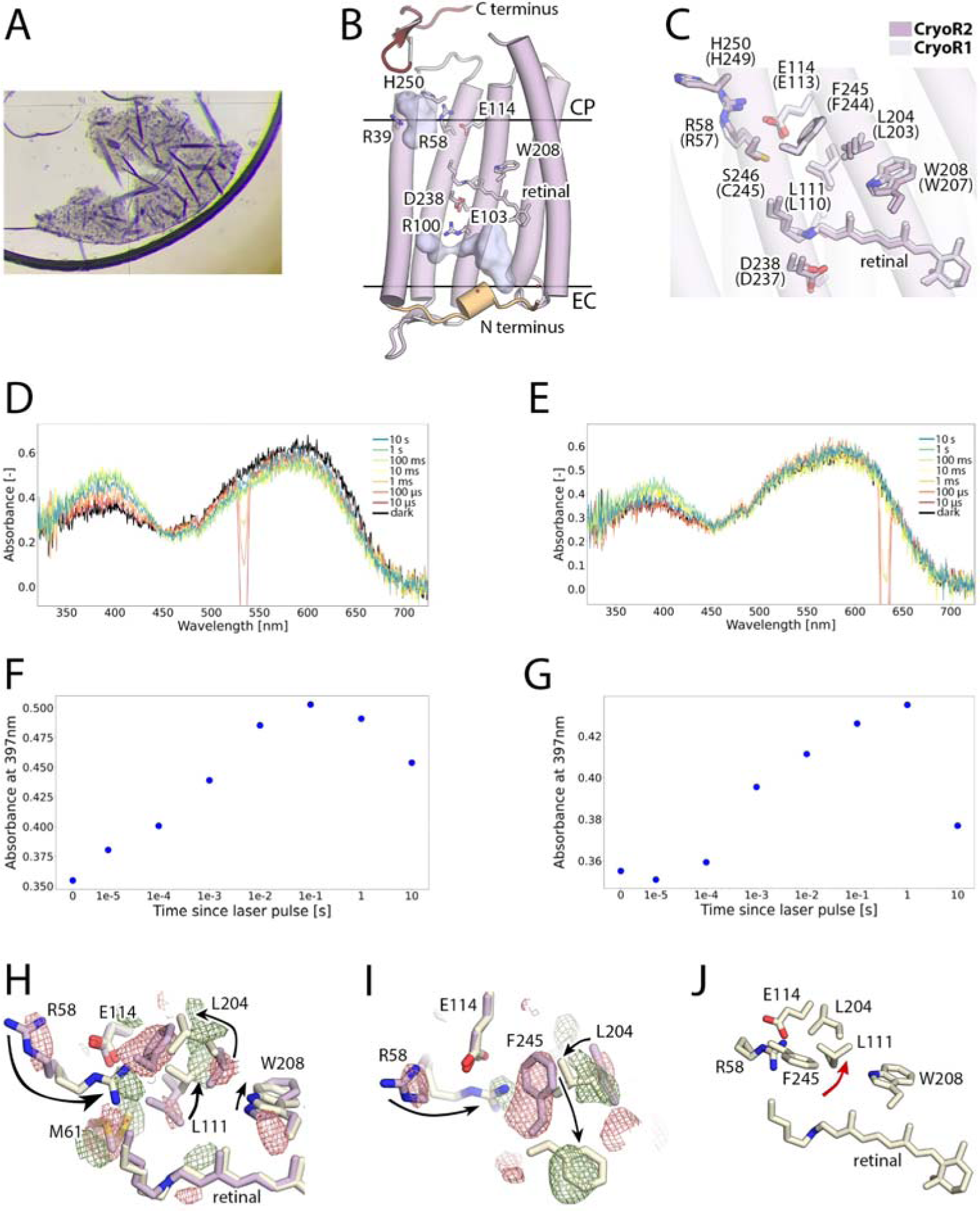
Crystal structures of CryoR2. **A.** Crystals of CryoR2 grown using *in meso* approach at pH 4.6. Crystals of two types (A and B) were found in the same crystallization well. **B.** Overall side view of the CryoR2 protomer at pH 4.6. N terminus is colored light orange. C terminus is colored red. **C.** Structural alignment of the CryoR1 (cryo-EM, pH 10.5, ground) and CryoR2 (X-ray, pH 4.6, ground) structures. Residues of CryoR2 are signed (the corresponding residues of CryoR1 are given in parenthesis). **D.** Spectra of CryoR2 crystal. **E.** Spectra of CryoR2 crystal. Black line corresponds to the dark spectra. Then, the evolution of the spectrum is shown in the time range of 10 µs to 10 s following the nanosecond green (532 nm, panel **D)** and red (630 nm, panel **E**) laser illumination of the crystal. **F.** Evolution of the absorption in the range of 392-402 nm corresponding to the blue-shifted state in the CryoR2 crystals following the nanosecond green (532 nm) laser illumination. **G.** Evolution of the absorption in the range of 392-402 nm corresponding to the blue-shifted state in the CryoR2 crystals following the nanosecond red (630 nm) laser illumination. **H.** Side view of the structural rearrangements in CryoR2 upon cryotrapping of the M_2_ state. **I.** View from the cytoplasmic side of the structural rearrangements in CryoR2 upon cryotrapping of the M_2_ state. Difference F_olight_-F_odark_ electron density maps are contoured at the level of 3.0σ. Black arrows indicate the key rearrangements associated with the M_2_ state formation. **J.** The cytoplasmic side of the CryoR2 protomer in the M_2_ state obtained with X-ray crystallography. Red arrow indicates the flipped side chain of L111, not found in the M_2_ state of CryoR1.

**Fig. S15.**
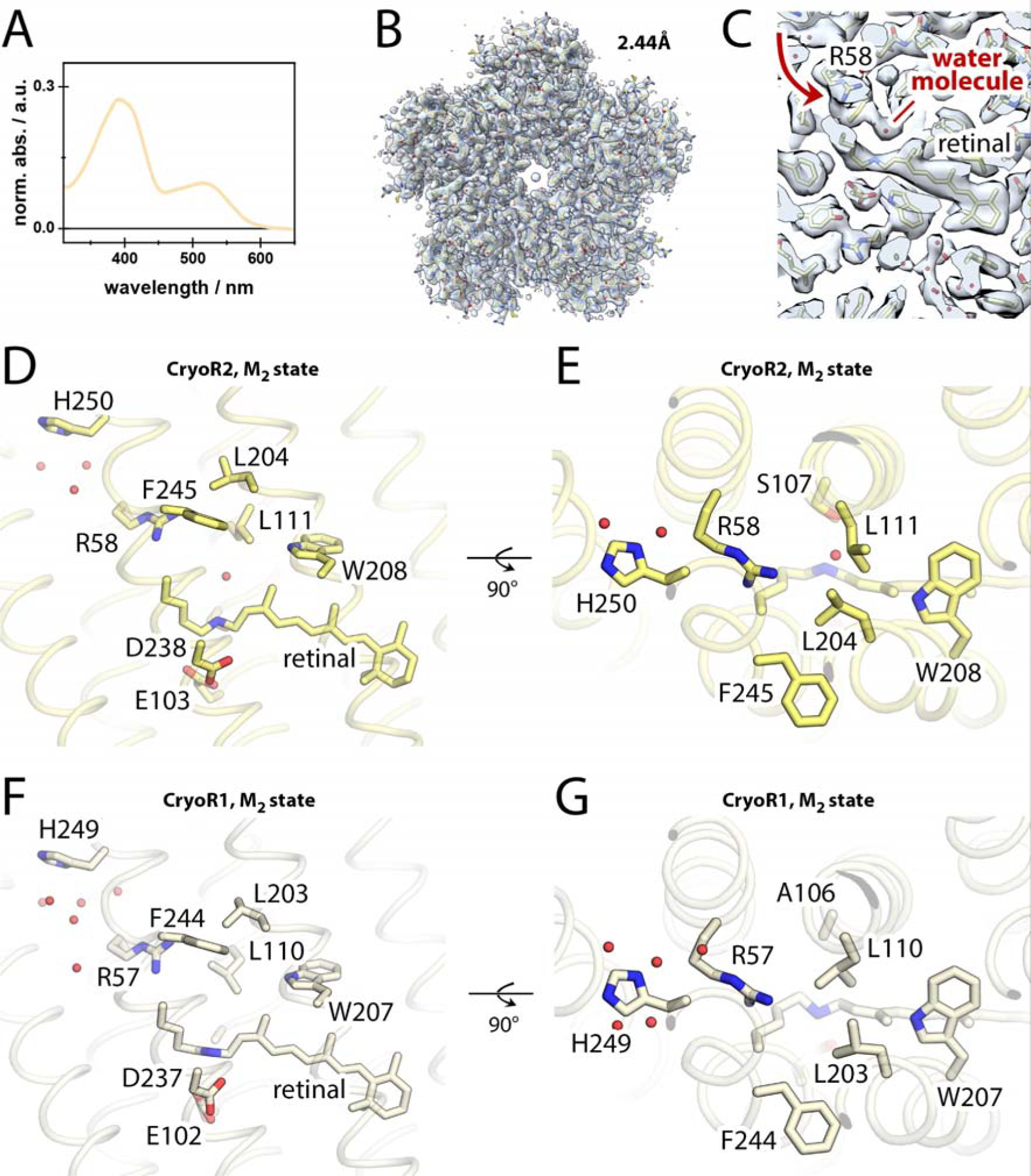
Cryo-EM structure of CryoR2. **A.** Spectra of the CryoR2 sample used for grid preparation. The sample was kept under overhead lights for 1h. **B.** Overall cryo-EM map of the pentameric CryoR2 at pH 8.0. **C.** Section view of the cryo-EM map in the region of retinal and R58. The flipped side chain of R58 in the M_2_ state is shown with a red arrow. An additional water molecule compared to the M_2_ state of CryoR1 is indicated with a red line. **D.** Side view of the cytoplasmic part of CryoR2 in the M_2_ state. **E.** View from the cytoplasmic side of the cytoplasmic part of CryoR2 in the M_2_ state. **F.** Side view of the cytoplasmic part of CryoR1 in the M_2_ state. **G.** View from the cytoplasmic side of the cytoplasmic part of CryoR2 in the M_2_ state.

**Fig. S16.**
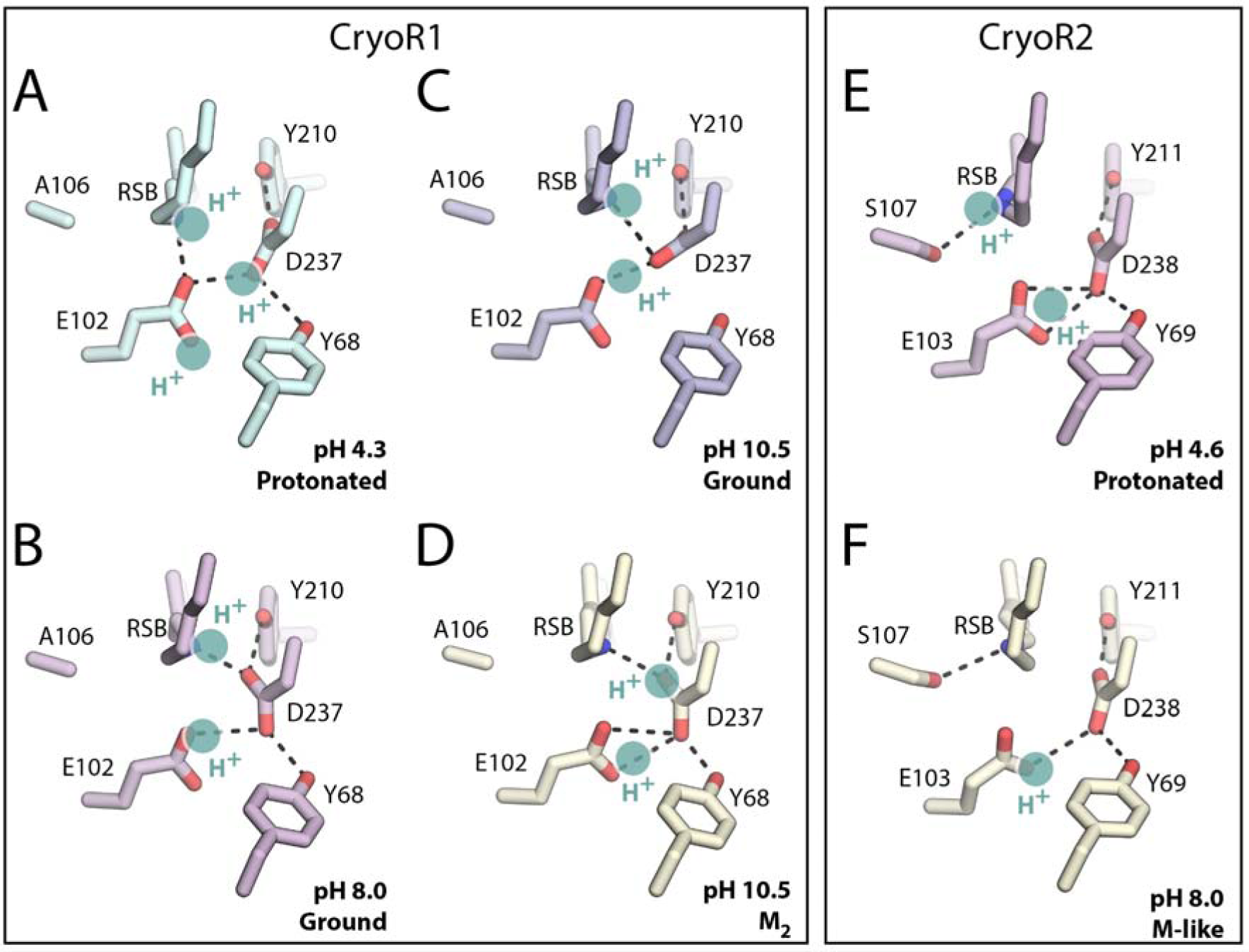
The RSB region of CryoR1 and CryoR2. **A.** The RSB region of CryoR1 at pH 4.3. **B.** The RSB region of CryoR1 at pH 8.0. **C.** The RSB region of CryoR1 at pH 10.5 in the ground state. **D.** The RSB region of CryoR1 at pH 10.5 in the M_2_ state. **E.** The RSB region of CryoR2 at pH 4.6. **F.** The RSB region of CryoR2 at pH 8.0. Putative protons in the RSB region are shown with cyan circles. H-bonds are shown with black dashed lines.

**Fig. S17.**
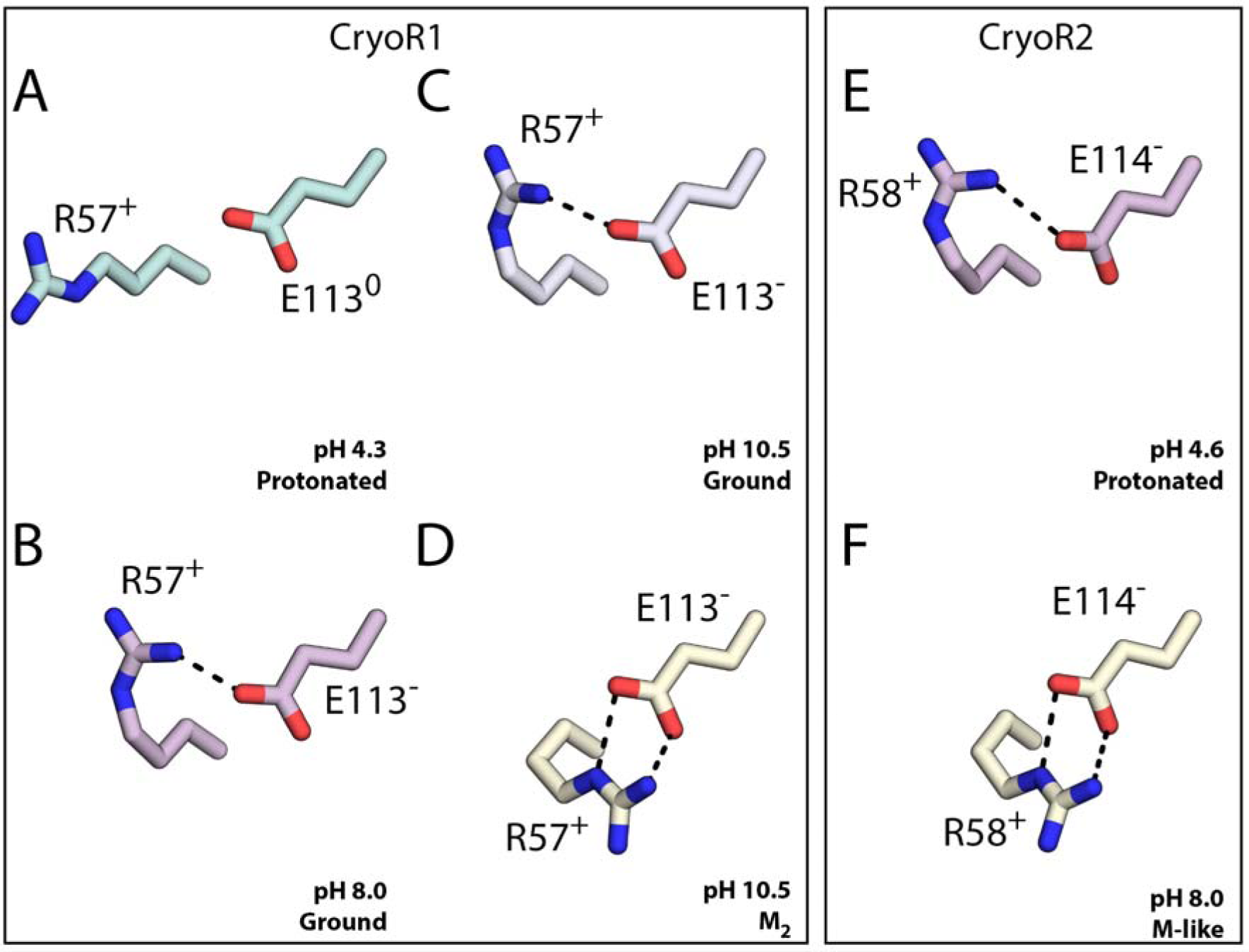
The arginine-glutamate pair at the cytoplasmic side of CryoRs. **A.** CryoR1 at pH 4.3. **B.** CryoR1 at pH 8.0. **C.** CryoR1 at pH 10.5 in the ground state. **D.** CryoR1 at pH 10.5 in the M_2_ state. **E.** CryoR2 at pH 4.6. **F.** CryoR2 at pH 8.0. H-bonds indicating the salt bridge between the residues are shown with black dashed lines.

**Fig. S18.**
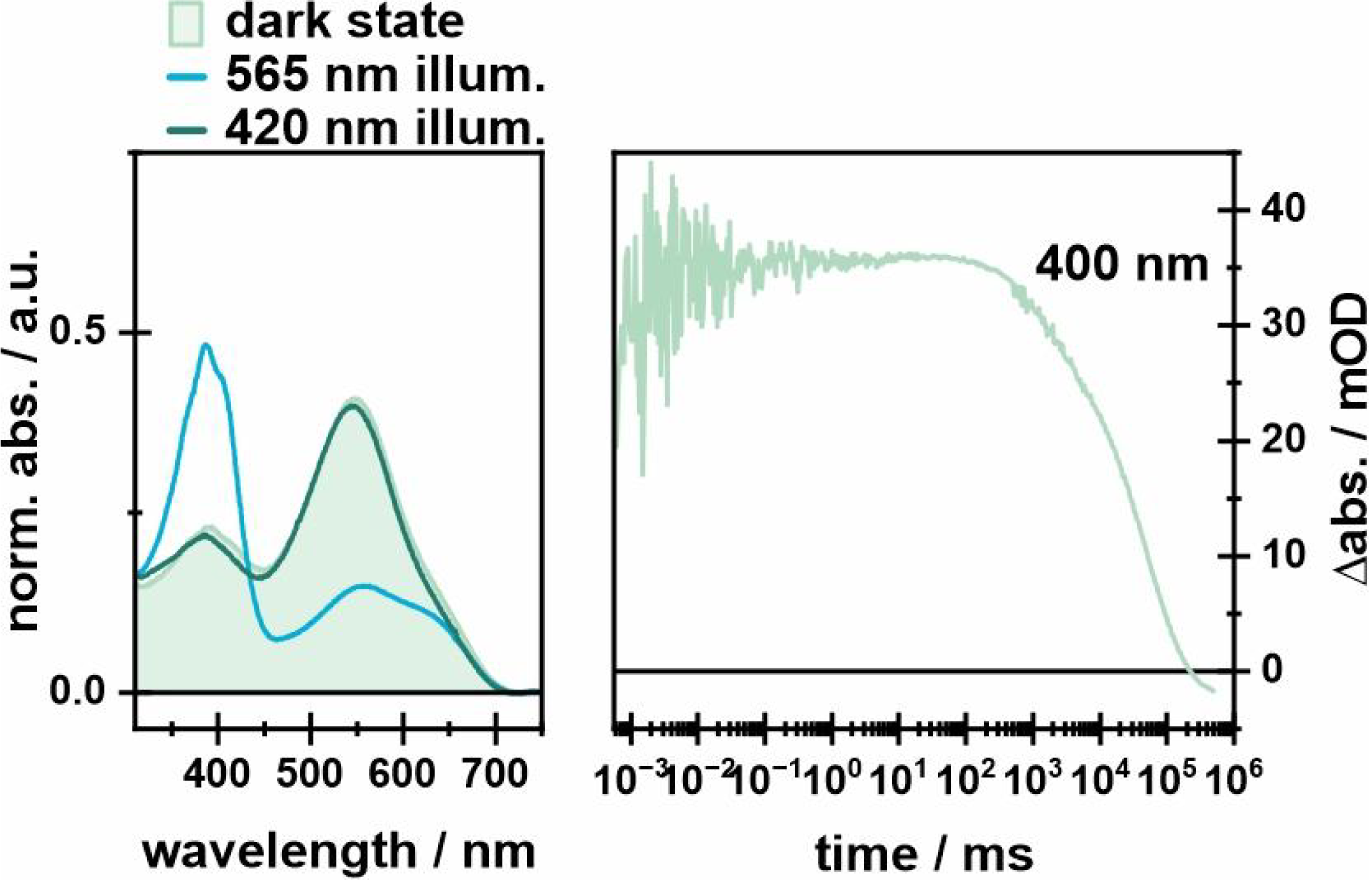
Illumination experiments of CryoR1 E113Q mutant. Left: Dark state and PSS spectra of CryoR1 E113Q mutant. First, the dark state spectrum was measured, followed by the PSS after illumination of the main absorption band for 100s and the illumination of the potentially rising band for 100s afterward. The 565 nm LED was operated at 100 mA and the 420 nm LED was operated at 10 mA. Additionally, the flash photolysis transient at 400 nm is shown to elucidate the slow kinetics of the M intermediate and the whole photocycle.

**Fig. S19.**
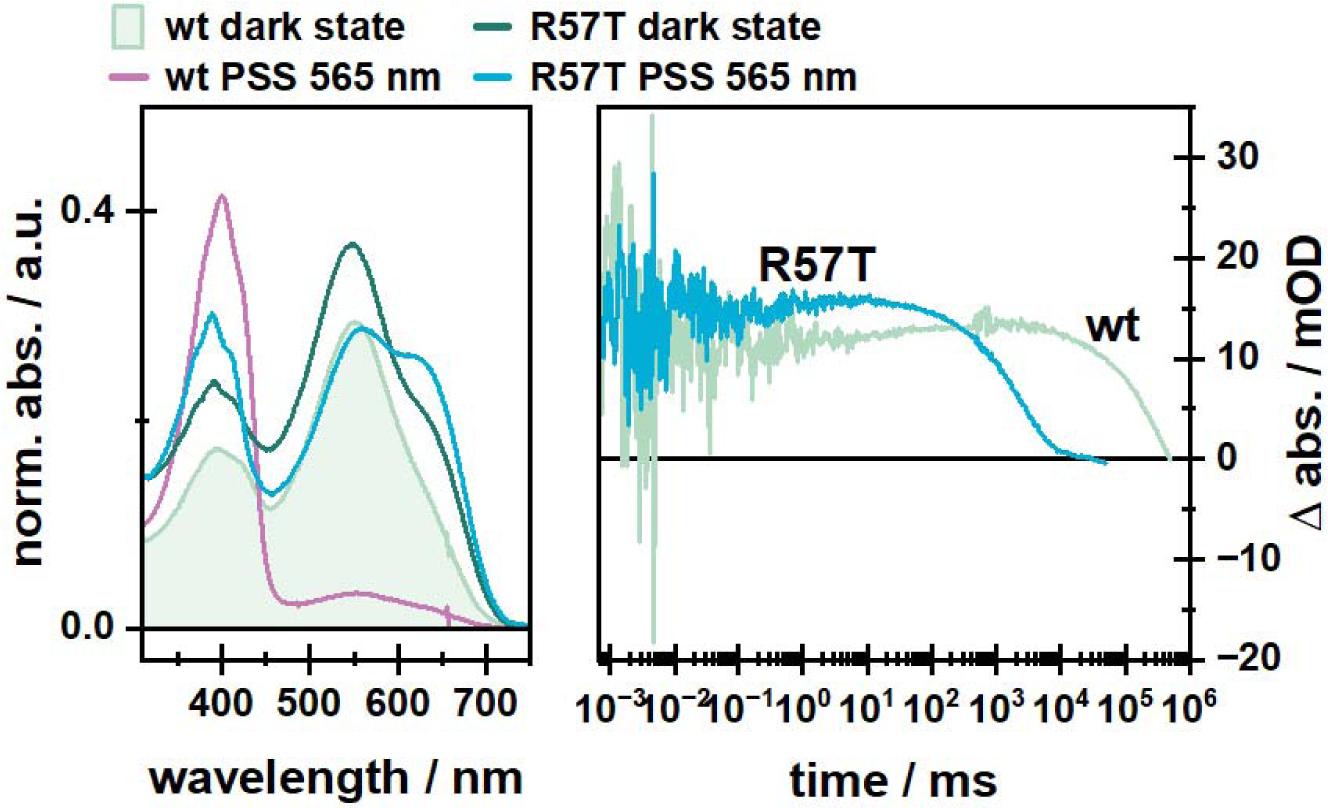
Comparison of spectroscopic properties of CryoR1-WT and R57T mutant. Dark state and PSS spectra of CryoR1-WT and R57T mutant. First of all, the dark state spectrum was measured, followed by the PSS after illumination of the main absorption band for 100s with a 565 nm LED operated at 100 mA. Additionally, the respective transient at 400 nm for both proteins is shown to illustrate the accelerated photocycle of the R57T mutant.

**Fig. S20.**
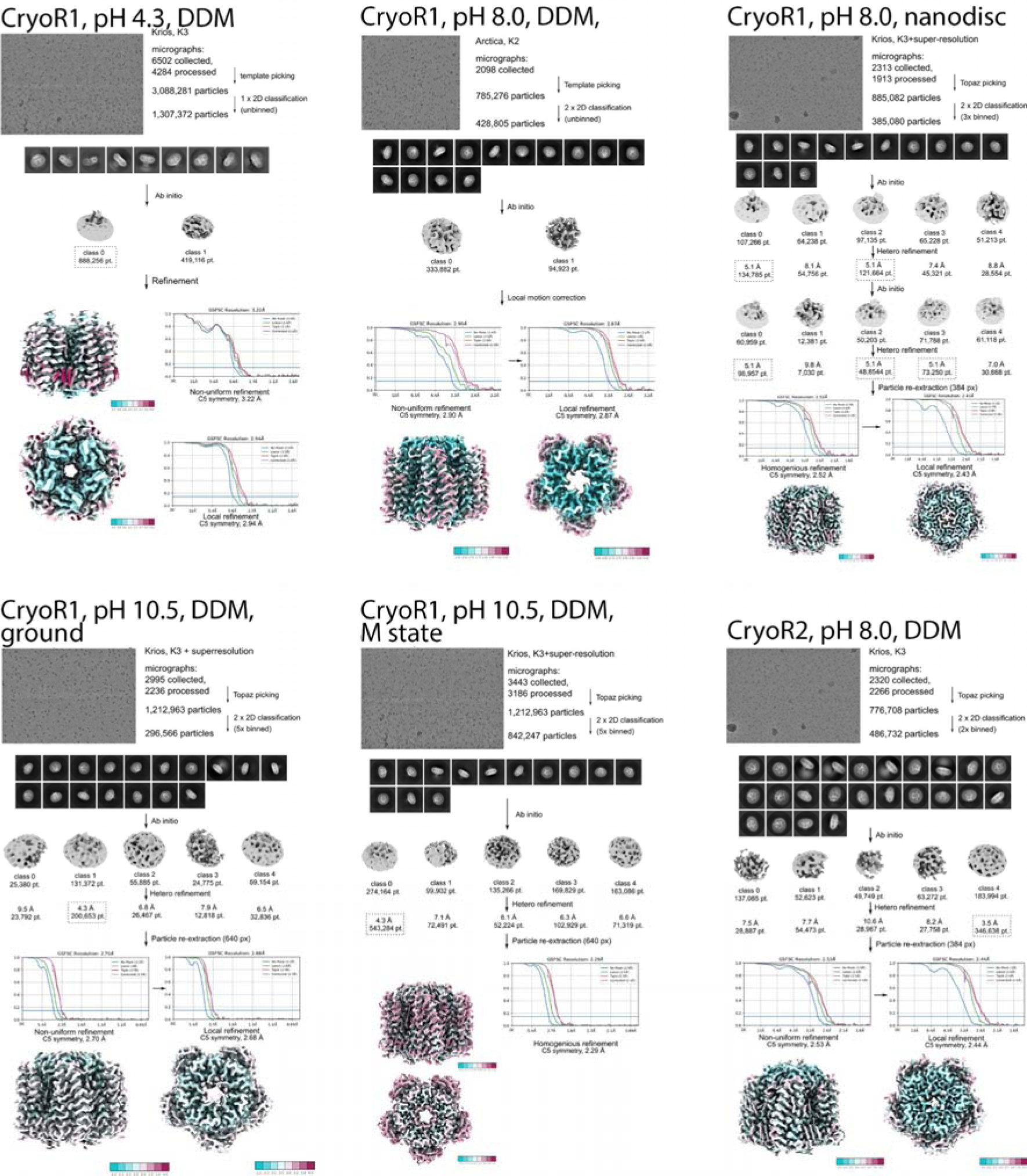
Workflow for solving the CryoR1 and CryoR2 structures using cryoSPARC. Overall resolution and local resolution are shown.

### Supplementary Tables

**Table S1.**
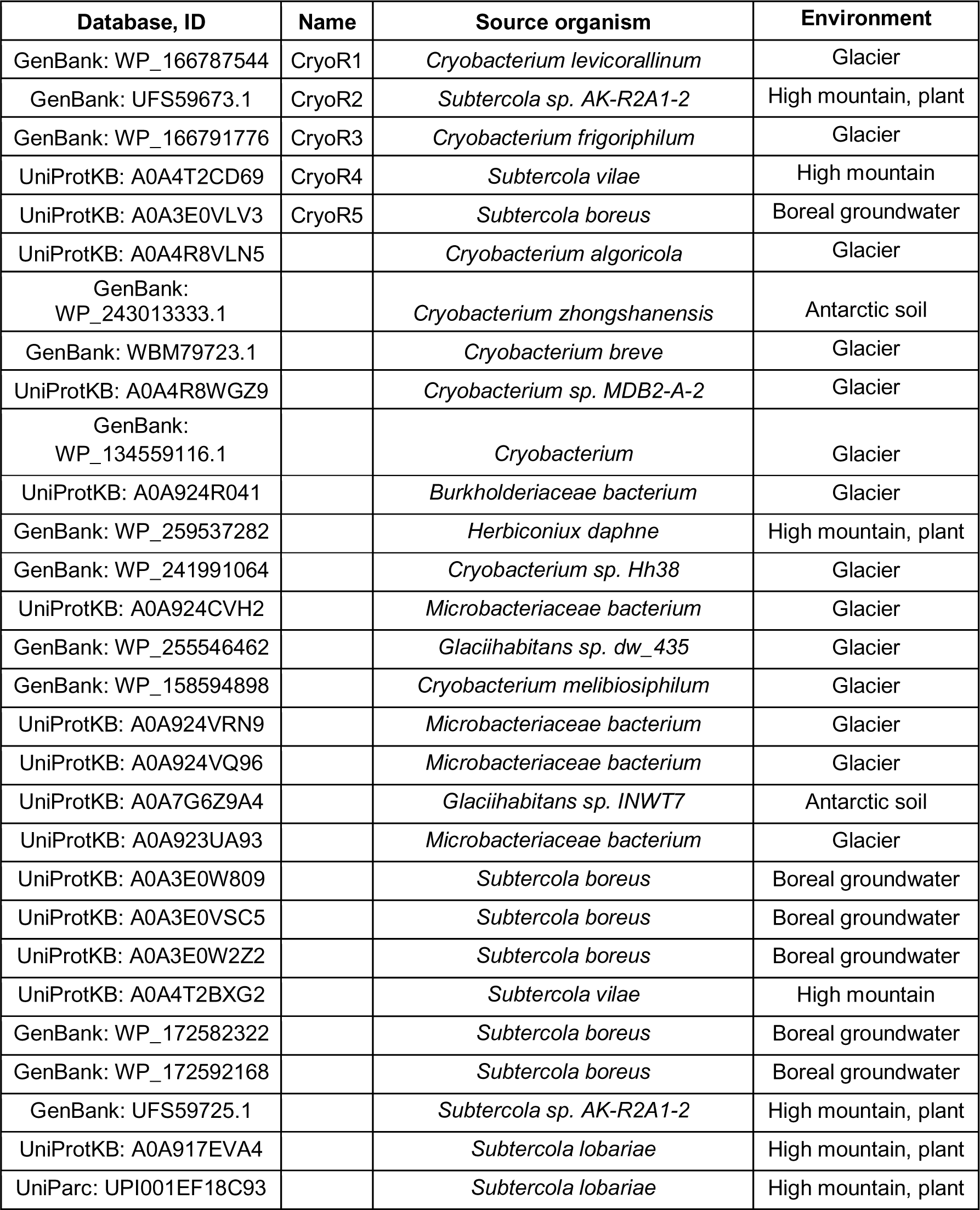
Source organisms of CryoRhodopsins.

**Table S2.** Summary of the structures of CryoRs obtained in the present work.

|  | **CryoR1** | **CryoR2** |
| --- | --- | --- |
| **acidic pH** | pH 4.3, cryo-EM, 2.94 Å, DDM | pH 5.2, X-ray crystallography, 2.65 Å, crystal |
| **neutral pH** | pH 8.0, cryo-EM, 2.4 3 Å, nanodisc | pH 8.0, cryo-EM, 2.44 Å, DDM |
|  | pH 8.0, cryo-EM, 2.9 Å, DDM |  |
| **alkaline pH** | pH 10.5, cryo-EM, 2.7 Å, DDM, dark (ground state) | - |
|  | pH 10.5, cryo-EM, 2.3 Å, DDM, illuminated (M<sub>2</sub> state) |  |

**Table S3.**
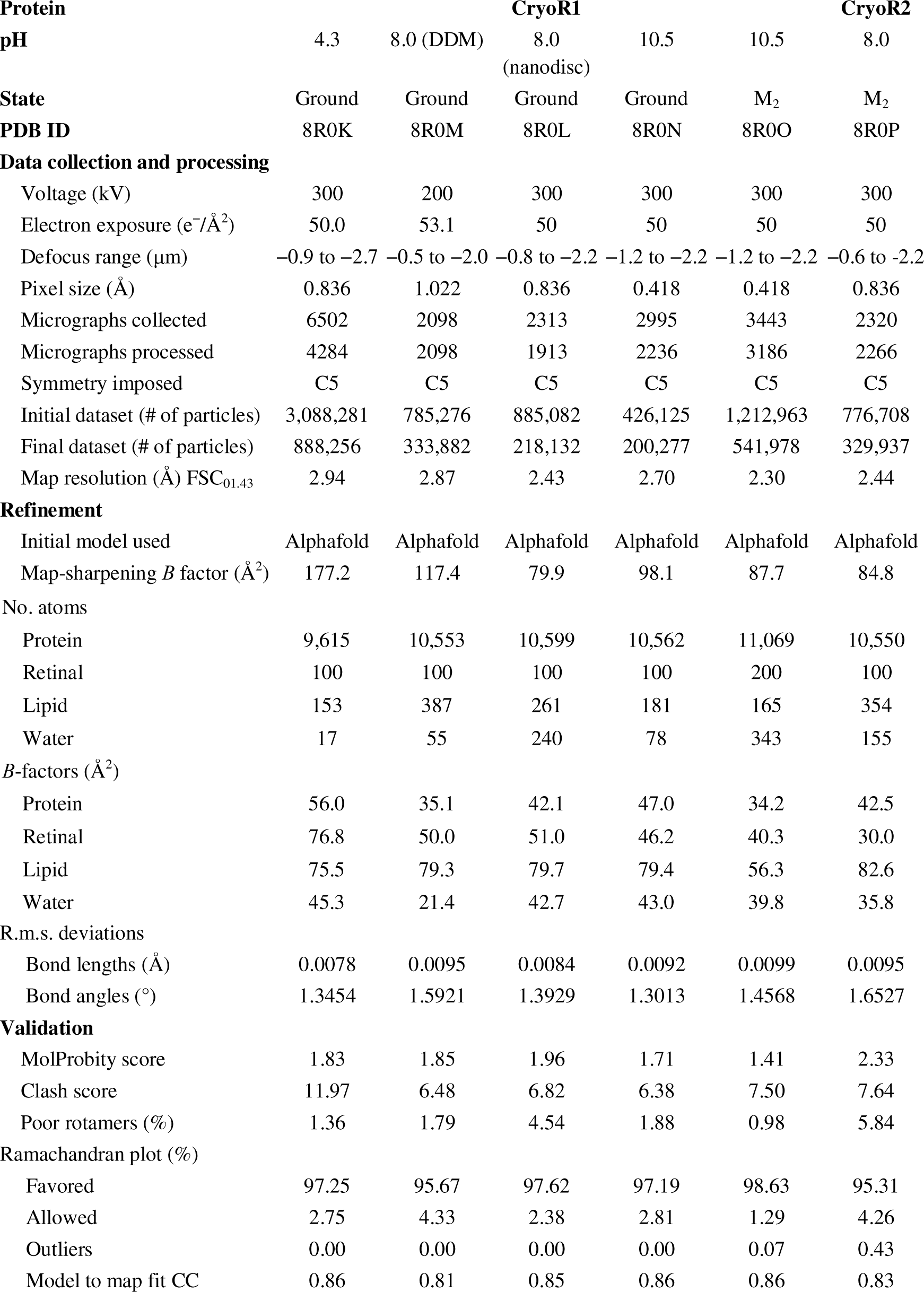
Cryo-EM data collection, refinement, and validation.

**Table S4.**
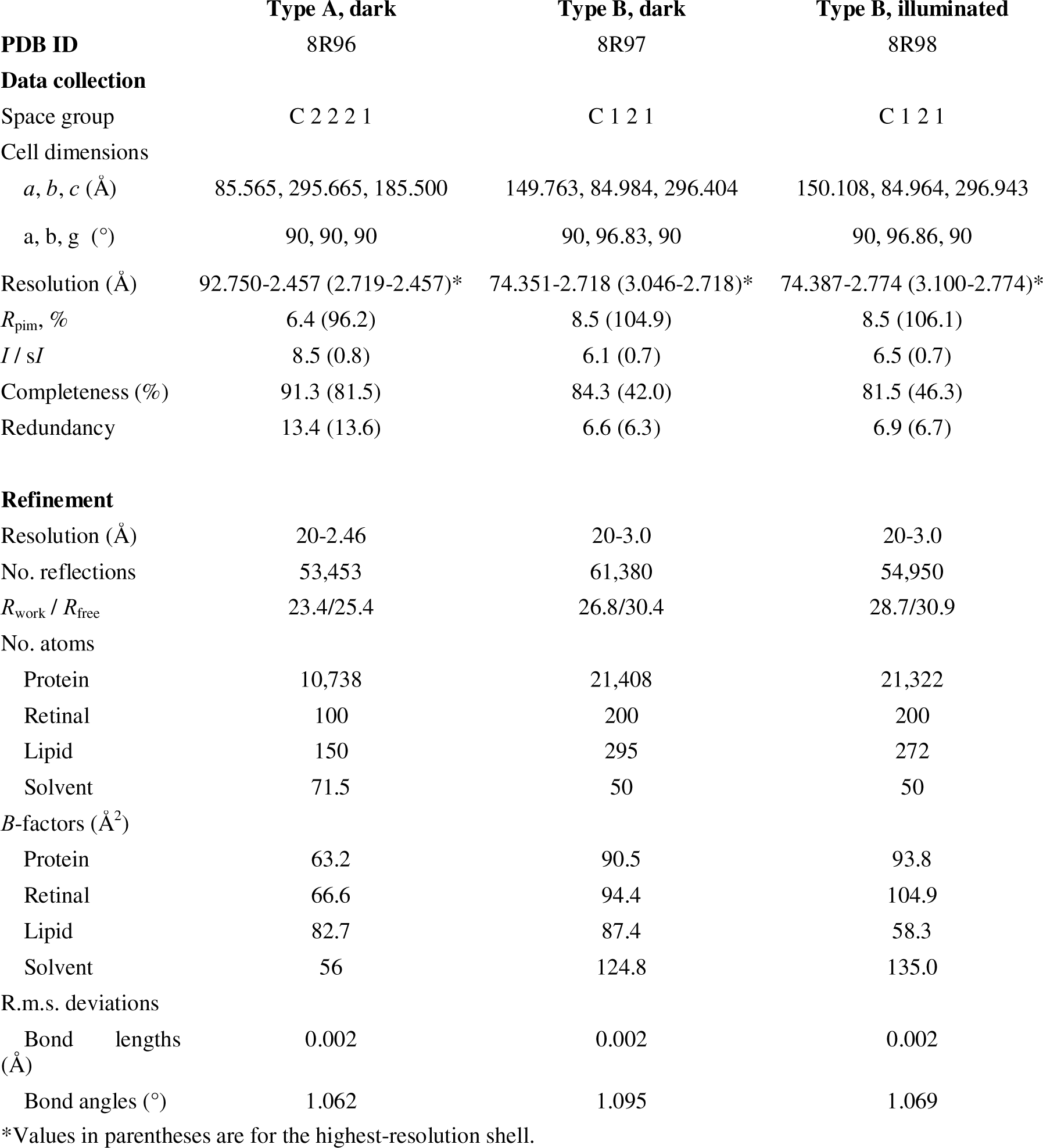
X-ray crystallography data collection and structure refinement statistics on CryoR2.

### Supplementary Note 1. Structural rearrangements in the M_2_ state of CryoR2

While for CryoR1 we were able to obtain the structures of the ground and M_2_ states individually under the same conditions, in the case of CryoR2 we obtained several structures of the protein under different conditions using different methods. Nevertheless, they provide essential information on the structural rearrangements associated with the M state formation in CryoR2.

First, we crystallized CryoR2 using *in meso* approach and obtained two types (A and B) of purple crystals at pH 4.6 (Fig. S14A). Using the crystals of type A (C2221) we obtained the structure of CryoR1 at 2.45 Å with X-ray crystallography. The crystals of type B (C2) diffracted more anisotropically to 3.0 Å and revealed a similar structure to that obtained with type A crystals. Therefore, we further discuss the 2.45 Å crystal structure of CryoR2.

The structure of CryoR2 shows similar overall organization to that of CryoR1 in the ground state (Fig. S14B,C). As in CryoR1, the extracellular side of CryoR2 has a large internal cavity, separated from the bulk by a short N-terminal α-helix (Fig. S14B). The side chain of R58 (R57 in CryoR1) is directed to the cytoplasmic bulk but not to the RSB. The residues in the helices F and G and the backbone of helix G are located same to the analog residues of CryoR1 in its ground state, including residues L204, W208, and F245 (L203, W207, and F244 in CryoR1) (Fig. S14C). Microspectrophotometry showed that a major fraction of CryoR2 molecules in crystal contains protonated RSB counterion complex as indicated by the red-shifted maximum absorption wavelength (Fig. S14D,E). A part of molecules retain the deprotonated complex (Fig. S14D,E). Lastly, a small fraction of the protein molecules is in the blue-shifted state characteristic for the deprotonated RSB (Fig. S14D,E). The relatively low resolution didn’t allow us to resolve multiple conformations of the RSB counterion complex associated with different protonation states. Nevertheless, the structural and spectroscopy data suggest that the above-described structure reflects that of the ground state of CryoR2.

Next, time-resolved spectroscopy on the CryoR2 crystal showed a pronounced formation of the blue-shifted state upon green (532 nm) or red (630 nm) light illumination (Fig. S14D-G). Noteworthy, the occupancy of the blue-shifted state was larger in the case of green light illumination, indicating that the red-shifted form with protonated counterion complex likely is not photoconvertible, similar to that observed for the red-shifted state of CryoR1 at pH 3.5. We cryotrapped the blue-shifted intermediate state in the crystal of CryoR2 using the protocol similar to that we used in recent works for the trapping of the M states of BR^43^ and *Bc*XeR^48^ (see Methods for details). Due to the limited amount of the type A crystals, we used type B crystals for the cryotrapping of the intermediate state. The F_olight_-F_odark_ difference electron density maps at 3.0 Å clearly showed the rearrangements at the cytoplasmic side of CryoR2, including the flip of the R58 side chain towards the RSB and synchronous reorientations of the residues in helices F and G similar to those found in the M_2_ state of CryoR1 (Fig. S14H,I). Different from CryoR1, the maps suggest the flip of the L111 residue (L110 in CryoR1) in the M_2_ state of CryoR2 (Fig. S14H,J). Since the cryotrapping protocol results in the accumulation of the longest-living intermediate, the observed rearrangements were assigned to the M_2_ state of the CryoR2 photocycle.

In addition to X-ray crystallography, we obtained a single-particle cryo-EM structure of CryoR2 at pH 8.0 at 2.44 Å resolution (Fig. S15A,B). The sample was kept under the overhead light before the application to the cryo-EM grid. Under such conditions, the blue-shifted M_2_ state is accumulated in the sample and is represented in the major fraction of protein molecules (Fig. S15C). The cryo-EM map revealed the crystal structure of the cryotrapped M_2_ state. Due to higher resolution, more details were resolved in the cryo-EM maps. Thus, the retinal is in likely the 13-*cis* conformation, similar to that of CryoR1 in the M_2_ state (Fig. S15B). The side chain of R58 is oriented towards the RSB (Fig. S15B,D,E). The L111 side chain is flipped compared to the ground state structure and an additional density is found near the RSB in the M_2_ state of CryoR2. We assigned it to a water molecule, which likely stabilizes the deprotonated form of the RSB in the intermediate. The RSB is additionally stabilized by S107 of the functional motif.

In summary, X-ray crystallography, cryo-EM, time-resolved absorption microspectrophotometry on crystals, and the cryotrapping of the intermediate state allowed us to show that the structural changes associated with the formation of the M_2_ state in CryoR2 are very similar those shown in the case of CryoR1. Since the arginine at the cytoplasmic side plays one of the key roles and is a unique characteristic residue in the entire CryoRs group, we suggest that the flipping motion of the arginine is conserved within all CryoRs.

At the same time, we found differences between the M state structures of CryoR1 and CryoR2. Namely, both cryo-EM and X-ray crystallography data show the flip of the L111 residue in the intermediate state of CryoR2, while no flip of L110 (analog of L111 in CryoR2) in the M_2_ state of CryoR1. Furthermore, while in CryoR1 the RSB is connected to the D237 and oriented towards helix G in both the ground and the M_2_ states, in CryoR2 the RSB is always oriented to helix C and is H-bonded to the S107. Thus, we speculate that the differences between the M_2_ state structures of two CryoRs originate from the more polar environment of the cytoplasmic side of the RSB region in CryoR2 than that of CryoR1. The role of the additional water molecule remains elusive; however, we speculate that it stabilizes the deprotonated RSB in the M_2_ state.

### Supplementary Note 2. Protonation states of the counterions in CryoRs

In the course of the study of CryoRs, we identified unusual spectral behavior of the members of this group. Together with the extremely long-living blue-shifted state, the RSB counterion complex of CryoR1 and CryoR2 demonstrate multiple protonation states as indicated by the pH titration of the maximum absorption spectra (Fig. S7). The structural data on CryoR1 at different pH values and in various functional states allowed us to get insights into the mechanisms of the proton storage at the RSB counterion complex of these rhodopsins.

In CryoR1, there are three major spectral states corresponding to the different protonation states of the RSB and its surroundings. According to the spectroscopy data, these states are nicely represented at pH ranges of 2.5-4.5 (620 nm), 7.5-9.0 (550 nm), and 10.5-11.5 (540 nm). To understand the differences of the maximum absorption wavelengths, we obtained the cryo-EM structures of CryoR1 at pH 4.3, 8.0, and 10.5 to have representative models for all three states identified in the spectroscopy experiments (Table S2).

Surprisingly, in the RSB region, the two counterions (E102 and D237) interact directly with each other in the ground state at all studied pH values (Fig. S16). This indicates that there is always at least one proton in the RSB counterions complex of CryoR1. In the 2.43 Å resolution structure of CryoR1 at pH 8.0, D237 is stabilized by Y68 and Y210 and interacts directly with the protonated RSB, which indicates that the residue is likely deprotonated (Fig. S16B). Thus, the proton is stored at E102 in the ground state of CryoR1 at neutral pH. At pH 10.5, the D237 side chain clearly reorients; however, we should note that at 2.7 Å resolution, it is impossible to univocally identify the exact position and orientation of the counterions. Nevertheless, in accordance with the spectroscopy data showing the ∼10 nm blue-shift of absorption maximum between pH 8.0 and 10.5 (Fig. S7), the conformation of the RSB region is also changed. The distance between E102 and D237 is shortened, which might be a signature of proton delocalization between the carboxylic residues (Fig. S16C). Thus, we suggest that the blue-shift of the spectrum upon pH increase from neutral to alkaline is associated with the proton redistribution rather than deprotonation of any of the counterions.

At pH 4.3, the conformation of the RSB-counterions complex is different from that at pH 8.0 and 10.5 (Fig. S16A). Namely, the protonated RSB is likely H-bonded to E102 instead of D237. D237 is H-bonded to the same oxygen atom OE1 of E102, but not to Y68, which tentatively indicates that D237 is protonated at pH 4.3. We speculate that E102 is also protonated; the proton is stored at the OE2 atom of the glutamic acid. The distance between E102 and D237 at low pH is shorter than that at pH 8.0, which might indicate proton delocalization between the residues, similar to that at pH 10.5. Indeed, at acidic pH values, CryoR1 demonstrates a notable (80 nm) red-shift of the absorption spectra (Fig. 3G; Fig. S7). Spectroscopy studies of the D237N, E102Q, and E102Q/D237N mutants, mimicking various protonation states of the RSB counterion complex, showed that none of these mutants result in such a strong red-shift as that found in the WT protein upon pH decrease (Fig. 3H,I). This supports our hypothesis on the proton delocalization with the unique resulting configuration of the RSB region, which is only possible within carboxylic, but not amide groups.

In the M_2_ state of CryoR1 at pH 10.5, the RSB is deprotonated; however, it is likely still connected to the D237 residue (Fig. S16D). Thus, we suggest that the RSB proton is transferred to D237 upon light illumination (Fig. 7A,B; Fig. S16C,D). E102 remains protonated in both the ground and the M_2_ states, as indicated by a direct H-bond between E102 and D237 similar to that in the ground state of CryoR1 at pH 8.0. Thus, both counterions are likely protonated in the M_2_ state. This observation is in line with the spectroscopy study of the E102Q/D237N mutant mimicking the fully protonated counterions complex (Fig. S8C). In the mutant, the RSB is deprotonated at both neutral and alkaline pH (Fig. S8C). This clearly indicates that the protonated counterions complex stabilizes the deprotonated form of the RSB, characteristic to the M_2_ state.

In summary, the counterion complex of CryoR1 is likely single-protonated at neutral and alkaline pH values with the proton localized at E102 but possibly delocalizing between E102 and D237 at alkaline pH. Single protonation allows an efficient proton transfer from the RSB to D237 with the rise of the blue-shifted state. The latter event results in the double protonated state of the E102-D237 pair, stabilizing the deprotonated RSB and thus contributing to the extremely slow decay of the M_2_ intermediate. At low pH, the RSB and both counterions are protonated, which hampers the formation of the blue-shifted state and significantly accelerates the photocycle (Fig. 3D,E; Fig. S5A). It should be noted, that at acidic pH values, the protonation of both counterions also likely hampers retinal isomerization as indicated by an extremely small fraction of the protein molecules entering the photocycle upon light illumination (Fig. 3E).

In the case of CryoR2, the structural data on various protonation states is limited. Nevertheless, both the cryo-EM and the crystal structures show that, in contrast to CryoR1, the RSB of CryoR2 is H-bonded to S107 (A106 in CryoR1) of the functional motif (Fig. S16E,F). D238 is H-bonded to Y69 and Y211 at pH 4.6 and 8.0, tentatively indicating that the residue is deprotonated in a wide range of pH values. E103 interacts directly with D238 in both structures; thus, E103 is likely protonated. Since the cryo-EM structure of CryoR2 at pH 8.0 likely represents the M_2_ state, we conclude that the counterions complex in the M_2_ intermediate of CryoR2 is single protonated, unlike that of CryoR1. This is supported by the spectroscopy analysis of the E103Q mutant of CryoR2, which showed similar spectral properties to those of E102Q/D237N of CryoR1 (Fig. S8C,D). Namely, the RSB is deprotonated in the E103Q mutant indicating that single protonation of E103 is sufficient to stabilize the M_2_ state of CryoR2. These results also likely mean the deprotonated RSB counterions complex at neutral and alkaline pH in the ground state of CryoR2.

At low pH, we have not observed any structural footprints of the D238 protonation. Similar to the M_2_ state at pH 8.0, the E103 residue is considered protonated as it forms an H-bond with deprotonated D238 (Fig. S16E). The deprotonation of the RSB proceeds effectively in CryoR2 at pH 3.5 (Fig. S5B); therefore, the RSB counterions complex is likely single protonated under these conditions in the ground state. Thus, we hypothesize that, unlike in CryoR1, the counterion complex of CryoR2 is deprotonated at neutral and alkaline pH, and is single protonated at low pH values (Fig. S16E,F). This results in the blue-shifted state formation at all studied proton concentrations, but also explains the differences in the spectral behavior of CryoR1 and CryoR2 (Fig. S2).

Thus, structural data suggest the alteration of the amino acid residue of the functional motif (A106/S107 in CryoR1/CryoR2) to be the key determinant of the spectral variations within the CryoRs group. At the same time, spectroscopy data on the CryoR1 and CryoR2 mutants together with high-resolution structures of the proteins at different pH values demonstrate that the proton is likely transferred from the RSB to the counterion complex and stored in it; the protonated complex stabilizes the deprotonated form of the RSB contributing to the extremely slow decay of the blue-shifted intermediate in CryoRs.

## Notes

### Competing Interest Statement

The authors have declared no competing interest.

### Summary of Updates

Section on the electrophysiology study of CryoR1 (Wavelength- and voltage-dependent proton translocation in CryoR1) was added to reveal ion translocation function of the rhodopsin. The Abstract, Introduction, and Discussion sections were revised in line with the observed ion transport activity.

